# Thermodynamic Coupling of the tandem RRM domains of hnRNP A1 underlie its Pleiotropic RNA Binding Functions

**DOI:** 10.1101/2023.08.17.553700

**Authors:** Jeffrey D. Levengood, Davit Potoyan, Srinivas Penumutchu, Abhishek Kumar, Yiqing Wang, Alexandar L. Hansen, Sebla Kutluay, Julien Roche, Blanton S. Tolbert

**Author notes:** Corresponding authors (J. Roche), (B. S. Tolbert).

## Abstract

The functional properties of RNA-binding proteins (RBPs) require allosteric regulation through inter-domain communication. Despite the foundational importance of allostery to biological regulation, almost no studies have been conducted to describe the biophysical nature by which inter-domain communication manifests in RBPs. Here, we show through high-pressure studies with hnRNP A1 that inter-domain communication is vital for the unique stability of its N-terminal domain containing a tandem of RNA Recognition Motifs (RRMs). Despite high sequence similarity and nearly identical tertiary structures, the two RRMs exhibit drastically different stability under pressure. RRM2 unfolds completely under high-pressure as an individual domain, but when appended to RRM1, it remains stable. Variants in which inter-domain communication is disrupted between the tandem RRMs show a large decrease in stability under pressure. Carrying these mutations over to the full-length protein for *in vivo* experiments revealed that the mutations affected the ability of the disordered C-terminus to engage in protein-protein interactions and more importantly, they also influenced the RNA binding capacity. Collectively, this work reveals that thermodynamic coupling between the tandem RRMs of hnRNP A1 accounts for its allosteric regulatory functions.

## Introduction

The heterogenous nuclear ribonucleoprotein A1 (hnRNP A1) is a ubiquitous RNA binding protein that regulates RNA metabolism both under normal and pathological cellular conditions, including viral infections [1-3]. As a general regulator of RNA biology, hnRNP A1 engages with transcripts from the moment they are synthesized, processed to maturity, exported from the nucleus, and translated into protein products [1, 4]. HnRNP A1 imparts its broad functions via a domain organization that consists of tandem RNA Recognition Motifs (RRMs), collectively known as UP1, at its N-terminus and a C-terminal low complexity domain (LCD_A1_) that is mostly disordered but engages in heterotypic protein-protein and protein-RNA interactions (**Fig. 1A**) [4-6]. Efforts to reconcile the pleiotropic functions of hnRNP A1 have led to multiple structures of its UP1 domain, each showing different mechanisms of RNA or DNA recognition [7-10]. These structures along with the broad RNA binding landscape of hnRNP A1 suggests that the protein binds its various targets via idiosyncratic mechanisms [1]. The underlying physicochemical properties by which hnRNP A1 achieves cognate RNA recognition remain incompletely understood, however. Adding to the hnRNP A1-RNA recognition paradox, its RRM domains are nearly identical both in sequence composition and three-dimensional structure, yet they exhibit contextual differences in RNA binding properties [9, 11, 12].

**Figure 1.**
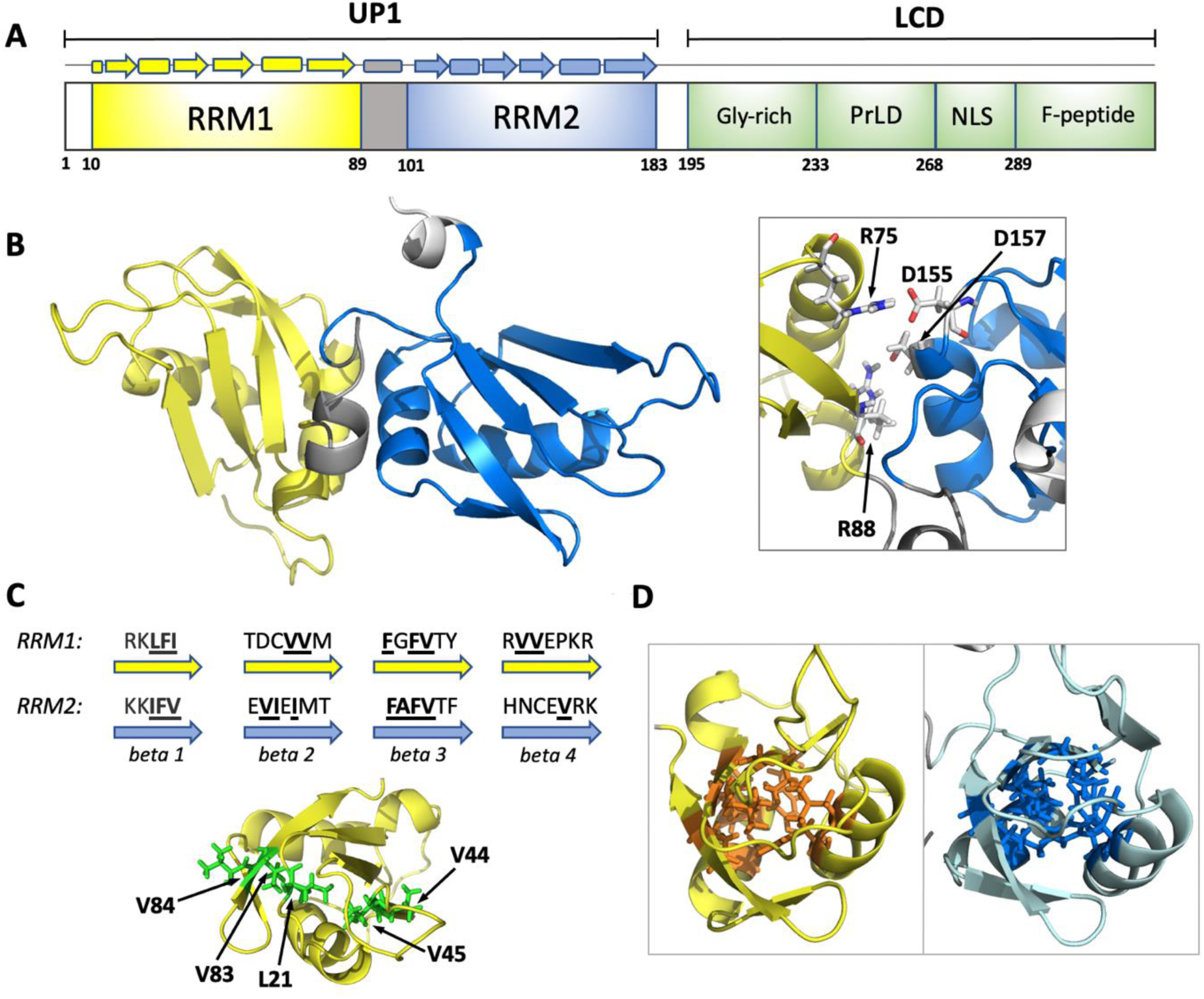
(**A**) Schematic representation of hnRNP A1-A sequence with the N-terminal Unwinding Protein-1 domain (UP1) encompassing two RNA Recognition Motifs (RRM1 and RRM2) and the C-terminal Low-Complexity Domain (LCD) encompassing the glycine-rich region (Gly-rich), prion-like domain (prLD), nuclear localization signal (NLS), and phosphopeptide (F-Peptide). (**B**) Three-dimensional structure of the UP1 domain (pdb 1U1R) showing RRM1 in yellow, RRM2 in blue, and the linker between the two RRMs in gray. Insert highlights the two salt-bridges (R75:D155 and R88:D157) at the interface between RRM1 and RRM2. (**C**) (*upper*) Sequence comparison of the four-stranded β-sheet found in RRM1 and RRM2. Non-polar amino acids composing the core of each RRM are shown in bold fonts. (l*ower*) Close-up of the hydrophobic patch present in the core of RRM1. (D) Close-up view of the RRMs highlighting the higher packing density of RRM1 core (left) compared to RRM2 (right).

A previous crystal structure from our lab demonstrated that hnRNP A1 can bind to RNA exclusively through its RRM1 domain and an inter-RRM linker to form complexes with 1:1 stoichiometries [7]. Although RRM2 did not contact RNA, we demonstrated that its physical coupling to RRM1, stabilized by two salt bridge interactions (R75:D155 and R88:D157) (**Fig. 1B**), was necessary to confer high-affinity RNA recognition to hnRNP A1. R75 and R88 are located on the alpha helical side at the inter-RRM interface. Mutating these two residues to Asp significantly reduced RNA binding affinity and was also accompanied by larger amplitude motions in the mutant protein indicating destabilization [7]. These results suggested that hnRNP A1 employs allosteric mechanisms to engage cognate RNA binding partners; however, the physicochemical basis of hnRNP A1 allostery was not obvious.

To explore the concept of hnRNP A1 allostery, we performed a series of ^15^N CPMG relaxation dispersion and high-pressure experiments (NMR and SAXS), complemented by extended molecular dynamic simulations and CLIP-seq to demonstrate that the physical coupling of the RRMs of hnRNP A1 impart uniform thermodynamic stability across the surface of the UP1 domain to allosterically control its RNA binding capacity. Analysis of the RRM1 and RRM2 sequences revealed differences in the β-sheet residues that serve as the hydrophobic cores of the RRM domains. Mutational transposition of these residues between RRM1 and RRM2 provided evidence for the strength of the hydrophobic cores being the source of domain stability. Notably, we found the core of RRM1 has a higher density of hydrophobic contacts than that of RRM2. Structure-based molecular dynamics simulations suggest that the difference in packing density is significant enough to give rise to differences in terms of thermodynamic stabilities. ^15^N-CPMG experiments validated that the packing density difference between RRM1 and RRM2 manifests with distinct conformational fluctuations, whereby RRM1 is essentially rigid in solution while RRM2 shows evidence of conformational exchange on a μs-ms time scale.

To gain atomic level insights into the differential stabilities of the RRMs, we used high-pressure solution NMR spectroscopy to probe the unfolding thermodynamics of the RRM domains, in isolation and in tandem within UP1. We found that the two RRMs display stark differences in stability under pressure despite the highly similar sequences giving rise to nearly identical structures. Indeed, RRM1 remains stable over a wide range of pressure while RRM2 is completely unfolded at 2.5 kbar. Studies conducted on the UP1 domain demonstrate that the inter-domain communication between RRM1 and RRM2 stabilize RRMs under high pressure conditions. Notably, we show that mutations designed to break inter-domain communication between the RRMs within UP1 or their structural transposition destabilize RRM2 without affecting the stability of the RRM1 domain.

Lastly, we examined the *in vivo* effects of allosteric regulation on RNA binding by performing CLIP-seq experiments with wild-type and the double salt bridge (R75D/R88D) variant of hnRNP A1 (A1^dm^). The consensus sequences identified by CLIP-seq for hnRNP A1 and the A1^dm^ variant were similar; however, comparative analysis of binding site occupancies across all transcripts immunoprecipitated show notable variations. For example, hnRNP A1 shows enriched binding to introns relative to the A1^dm^ mutant, which correlates with statistically significant changes in the RNAs bound by the two proteins. Importantly, the difference in binding capacity between hnRNP A1 and the A1^dm^ mutant correlates with the latter being defective in its ability to dimerize on the immunoprecipitated transcripts. When interpreted collectively, this study demonstrates that hnRNP A1 is an allosterically regulated RNA binding protein and that the physical coupling of its tandem RRMs is necessary to confer functional recognition of cognate RNA molecules. Perturbations that change the degree of inter-domain communication such as post-translational modifications or naturally occurring mutations would in turn influence the pool of RNAs regulated by hnRNP A1.

## Results

### Sequence and structural determinants of hnRNP A1 allostery

Since determinants of hnRNP A1 allostery would be reflected in its domain composition, we proceeded to compare the physicochemical properties of RRM1 and RRM2. The tandem RRMs of hnRNP A1 adopt identical folds composed of a four-stranded β-sheet and two α-helices (**Fig. 1 A,B**) [1, 2]. High resolution x-ray structures indicate that the tertiary structure of RRM1 is remarkably similar to that of RRM2 (Cα RMSD ∼ 0.95 Å, PDB 1U1R) [13]. Yet close examination of the amino acids composing the four antiparallel β-strands reveals interesting differences between RRM1 and RRM2. For example, β2 and β4 of RRM1 each contains two successive valines that are not present in RRM2 (V44, V45, V83, and V84). These valines, together with L21 located in the loop that connects β1 and β2, form a long hydrophobic patch that is unique to RRM1 (**Fig. 1C**). Valines from RRM1 are indeed replaced in RRM2 by an isoleucine and three polar and charged residues. The connecting loop also has one less hydrophobic residue, practically eliminating the hydrophobic patch.

Next, we compared the packing density of RRM1 and RRM2 by analyzing inter-side chain distances between residues composing their structural cores. We defined as structural core the 10 residues with lowest accessible surface area (ASA) in each RRM: L16, I18, L21, F34, G58, V60, Y62, V68, A71, and P86 for RRM1, and I107, V109, L121, F125, A149, V151, F153, V159, I162 and V177 for RRM2. We found that the average distance between the nearest side chain atoms within these core residues to be 2.8 Å for RRM1 and 3.1 Å for RRM2, suggesting that the core of RRM1 is slightly more densely packed than that of RRM2 (**Fig. 1D**). To gauge whether this slight packing difference could potentially result in a difference in thermodynamic stability, we conducted a set of molecular dynamics simulations using a structure-based potential (Go-model). We ran a series of all-atom simulations at various temperatures using a native-based Go-type potential energy function that portrays a perfect funneled energy landscape [14, 15]. By computing the change in specific heat capacity as a function of temperature, we found that the isolated RRM2 (residues 95-196) has a lower folding temperature compared to the isolated RRM1 (residues 1- 105), suggesting that RRM2 may potentially be thermodynamically less stable than RRM1 (**Fig. S1**).

### Evidence of differential μs-ms conformational dynamics of the RRM domains of hnRNP A1

To evaluate if the compositional differences between RRM1 and RRM2 manifest as unique physicochemical properties, we performed backbone ^15^N Carr–Purcell–Meiboom–Gill (CPMG) relaxation dispersion experiments. These experiments provide site-specific information on the contribution of microsecond to millisecond dynamic processes to the effective transverse relaxation rate constant (*R*_2_,_eff_ = *R*_2_° + *R*_ex_) [16, 17]. We conducted ^15^N-CPMG experiments on the UP1 domain (residues 1-196), isolated RRM1 (residues 1-105), and isolated RRM2 (residues 95- 196) (**Fig. S2**). Analysis of relaxation dispersion profiles collected for UP1 shows that the intact domain is rigid in solution with very few residues experiencing conformational dynamics on a μs-ms time scale (**Fig. S3**). A similar picture emerged from the analysis of the ^15^N-CPMG data collected for the isolated RRM1 for which only 9 amide groups appear to experience conformational dynamics. The magnitude of R_ex_ measured for these residues is small (< 4 s^-1^), indicating that the isolated RRM1 is predominantly rigid in solution (**Fig. 2A-C**). Of significance, we observed contributions to transverse relaxation for 19 residues of the isolated RRM2, with R_ex_ values 3-4 times larger than those measured for RRM1 (**Fig. 2A-C**). Residues of RRM2 experiencing the largest contribution from conformational exchange are located within β2 and between α2 and β4 (**Fig. 2D**).

**Figure 2.**
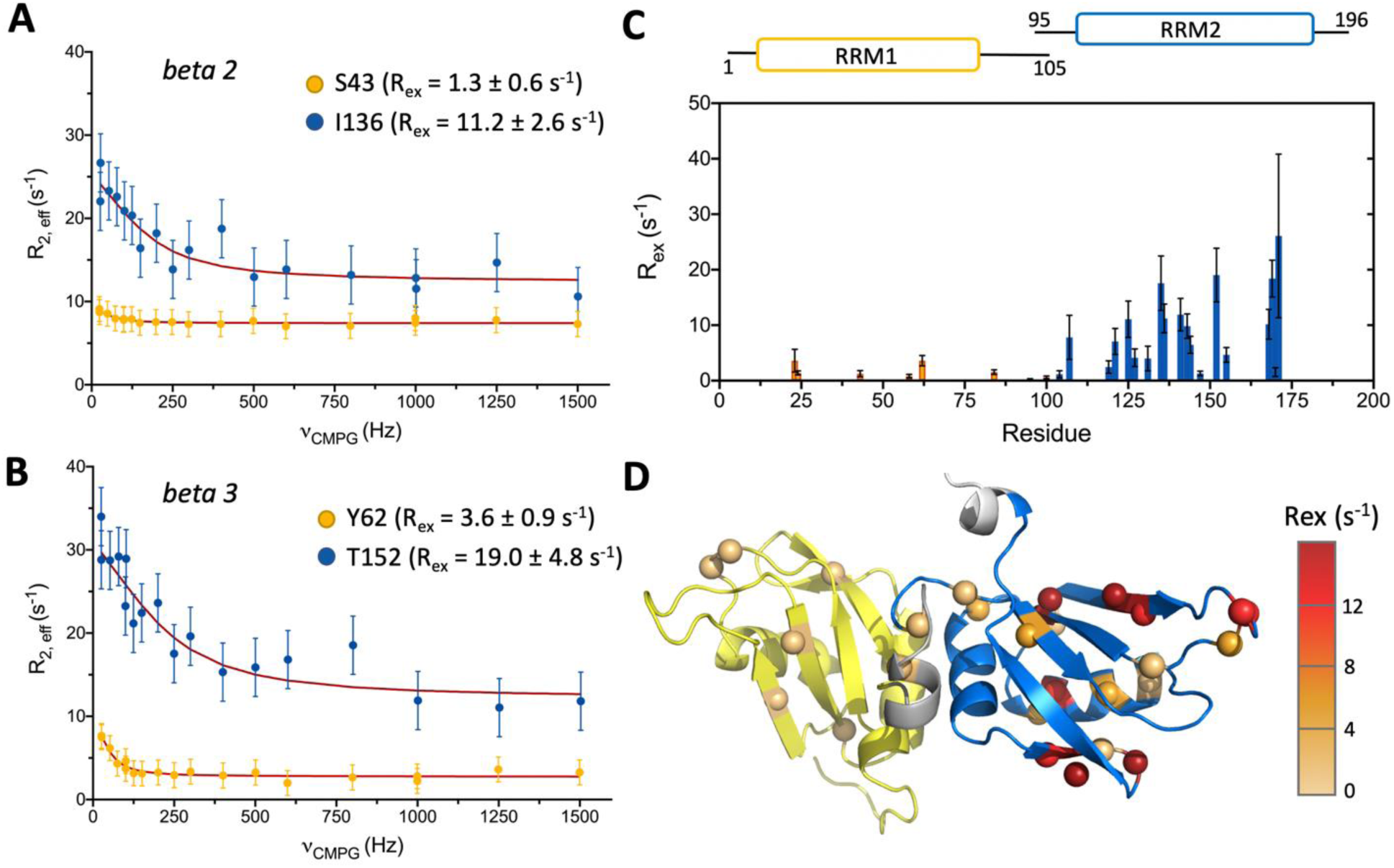
**(A, B)** Representative ^15^N relaxation dispersion profiles measured for residues within (**A**) the second and (**B**) third β-strand of the RRMs. Residues within RRM1 (S43 and Y62) are shown in yellow while residues within RRM2 (I136 and T152) are shown in blue. (**C**) Chemical exchange contribution to the ^15^N transverse relaxation rate (R_ex_) measured for amide resonances of the isolated RRM1 (yellow) and isolated RRM2 (blue). (**D**) R_ex_ values measured on the isolated RRMs are mapped on the structure of UP1, showing that residues experiencing significant conformational exchange are predominantly localized within RRM2 (blue). One unassigned RRM1 peak had R_ex_=1.5 ± 0.43

Overall, these experiments indicate that the isolated RRMs are exhibiting significantly different conformational dynamics in solution; RRM1 being essentially rigid while RRM2 shows evidence of conformational exchange on a μs-ms time scale. Connecting RRM1 and RRM2 via stabilization of the inter-domain interface in the context of UP1 abates the conformational dynamics of RRM2 to the level of RRM1.

### Pressure-induced unfolding reveals that the coupling of inter-RRM interface imparts uniform thermodynamic stability

Next, we probed the thermodynamic stability of several variants of the UP1 domain designed to evaluate the integrity of the inter-RRM coupling. When pressurized, protein NMR spectra typically display two types of perturbations: chemical shift changes, which are due to changes in protein surface-water interface and/or small compression of protein native conformations [18], and crosspeak intensity changes. Loss of crosspeak intensity as a function of pressure points to major conformational transitions on a slow time scale (relative to NMR time scale) such as folding/unfolding [19]. Here we monitored the pressure-induced changes of crosspeak intensity of UP1 variants by recording series of 2D ^1^H-^15^N HSQC experiments from 1 bar to 2.5 kbar.

Remarkably, our experiments show that the UP1 domain exhibit only minor changes in crosspeak intensity as a function of pressure (∼15% loss at 2.5 kbar) suggesting that UP1 remains predominantly native at high-pressure (**Fig. S4 A,B** and **Fig. 3A**). Similarly, we observed no major change for the isolated RRM1 with only ∼8% loss at 2.5 kbar, which indicates that RRM1 is thermodynamically stable under pressure (**Fig. S4 C,D** and **Fig. 3B**, yellow lines). By contrast, the isolated RRM2 displays a complete loss of native crosspeaks within 2.5 kbar (**Fig. S4 E,F** and **Fig. 3B**, blue lines). Decrease of native crosspeak intensity is accompanied by the appearance of new sets of crosspeaks with a narrow ^1^H chemical shift dispersion that is typically observed for protein unfolded conformations (**Fig. S4F**). Comparison of the native and unfolded crosspeak intensity as a function of pressure indicates that the conformational transition experienced by RRM2 is characteristic of a two-state unfolding transition with a mid-point p_1/2_ = ∼ 1.5 kbar (**Fig. S5**). This transition is fully reversible with no sign of protein precipitation.

**Figure 3.**
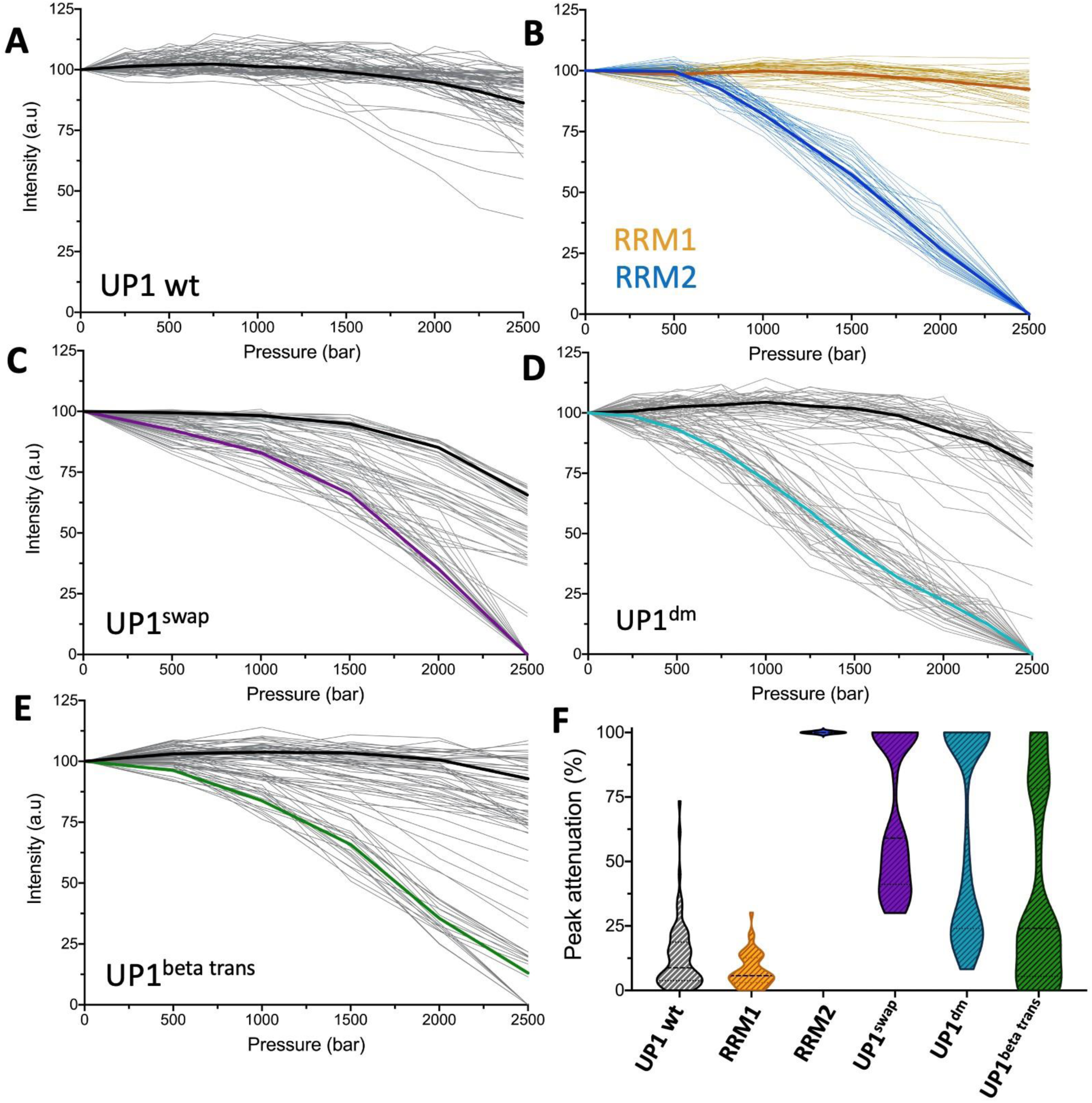
(**A-E**) NMR peak intensity profiles measured as a function of pressure for individual amide resonances of (**A**) wt UP1, (**B**) isolated RRM1 (yellow) and isolated RRM2 (blue), (**C**) UP1^swap^, (**D**) UP1^dm,^ and (**E**) UP1^beta^ ^trans^. The average peak intensity profile is shown for each variant with a bold line. (**F**) Distribution of the percentage of peak attenuation (based on the individual NMR peak intensities measured at 1 bar and at 2.5 kbar) for the amide resonances of wt UP1, isolated RRMs, and UP1 variants.

These three sets of experiments demonstrate that: (*i*) despite their structural similarities, the isolated RRM2 is significantly less stable under pressure than the isolated RRM1, and (*ii*) RRM2 is stabilized when linked to RRM1 in the full UP1 domain. To investigate the origin of the stabilization of RRM2 in the UP1 domain, we first analyzed a variant for which the position of the RRMs are swapped (UP1^swap^, **Fig. S2**). Prediction by Alphafold 2 [20] (shows that the interface between the two RRMs and their relative orientation are significantly modified in UP1^swap^ compared to the wild type UP1 (**Fig. S6**). When pressurized, UP1^swap^ appears to be much less stable than wt UP1 (**Fig. S7 A,B**). Interestingly, analysis of intensity profiles reveals two types of pressure sensitivity: a group of residues located within RRM2 that shows an almost complete loss of intensity at 2.5 kbar, and a group of residues mainly located within RRM1 that exhibits only minor changes as a function of pressure (**Fig. 3C**).

The differential pressure-induced unfolding of UP1^swap^ and UP1 indicates that the integrity of the inter-RRM interface is a determinant of the thermodynamic stability and coupling of tandem RRMs of hnRNP A1. As a test of this concept, we examined the stability under pressure of a double salt bridge (R75D/R88D) variant of UP1 (UP1^dm^). This variant was designed to disrupt the salt bridges (R75:D155 and R88:D157) found at the interface between the two RRMs (insert of **Fig. 1B**). We previously showed that UP1^dm^ binds cognate RNA substrates with significantly weaker affinity compared to wild type UP1 and that the salt bridge interactions stabilize the relative orientation of the tandem RRMs during the course of a short 10 ns molecular dynamic simulation [7]. Notably, figure 3D shows a similar clustering between pressure-sensitive and pressure-resistant residues for UP1^dm^ as observed for UP1^swap^. Two distinct groups of residues were also observed for UP1^dm^ based on intensity profiles: a group that shows almost complete intensity loss at 2.5 kbar (located within RRM2), and a group that displays only minor effect of pressure (mainly located within RRM1) (**Fig. S7 C,D** and **Fig. 3D**). These results demonstrate that the integrity of the interface between the tandem RRMs plays a crucial role in stabilizing RRM2 in the context of the UP1 domain. Modifying the interface, as in UP1^swap^, or disrupting key interactions between the RRMs, as in UP1^dm^, disrupts the thermodynamic (allosteric) coupling between RRM1 and RRM2 (**Fig. 3F**).

Finally, we sought to determine what factors contribute to the intrinsic lower thermodynamic stability of RRM2 compared to RRM1. We designed a variant of UP1 for which the sequence of RRM1 beta strands are replaced by those of RRM2 (and vice versa) while keeping the interface between the two RRMs mostly intact (UP1^beta-trans^, **Fig. S2**). We observed for this variant a group of residues with high sensitivity to pressure, mainly corresponding to core residues within the modified RRM1, while a second group of residues, mainly within the modified core of RRM2, show a much lower sensitivity to pressure (**Fig. S 7E,F**, **Fig. 3 E,F**). As described above, we identified both the extent of hydrophobic patches and packing density as the main structural differences between RRM1 and RRM2. Notably, the present results suggest that modifying these structural properties by transposing the sequences of β1-4 reverses the relative thermodynamic stability of RRM1 and RRM2.

### High-pressure SAXS experiments of intact hnRNP A1 suggests that the tandem RRMs influences the low complexity CTD

High pressure small angle x-ray scattering (SAXS) experiments were carried out with four variants of hnRNP A1 to complement the high-pressure NMR experiments described above: wt UP1, UP1^dm^ (**Fig. S2**), wt full-length hnRNP A1, and full-length hnRNP A1 R75D/R88D (A1^dm^). Each variant was tested at separate pressures from 0 to 2.5 kbar, in increments of 500 bar. The radius of gyration (Rg) was monitored for each variant at each pressure (**Fig. S8**). Results for wt UP1 analysis of radius of gyration (Rg) values confirmed that the protein is stable and doesn’t unfold under pressure. The Rg value measured for UP1^dm^ at atmospheric pressure is identical to that of the wt UP1, indicating no effect from the loss of the salt bridges on the overall globular structure of the protein. Yet a sharp increase in Rg was observed under pressure for UP1^dm^, confirming, in very good agreement with the NMR data, the loss of stability due to the loss of the inter-RRM salt bridges.

Next, we examined the effect of the double mutation R75D/R88D on the pressure stability of the full-length hnRNP A1. We observed a slight increase in Rg values under pressure for wt hnRNP A1, likely due to low complexity C-terminal domain elongating from a compact to an extended conformation, as recently reported [21]. For A1^dm^, the Rg measured at atmospheric pressure was found to be significantly larger than that of the wt hnRNP A1. A similar result was also observed by SEC-SAXS (not shown). As the UP1 domains are identical in size, such increase is likely due to an elongation of the C-terminus that is affected by the loss of the R75:D155 and R88:D157 salt bridge interactions that stabilize the inter-RRM interface in the UP1 domain. Application of pressure reveals a further increase in Rg values measured for A1^dm^, suggesting a complete unfolding of the UP1 domain (**Fig. S8**). These results confirm that the salt-bridge disrupting mutations dramatically affect the stability under pressure of the tandem RRMs, for both the full-length hnRNP A1 and isolated UP1.

### Molecular Dynamic simulations reveals that the integrity of the tandem RRMs impose long-range effects on the low complexity CTD

Molecular dynamic simulations were then performed to assess how the domains of hnRNP A1 interact with each other and how these interactions are disrupted by the R75D/R88D mutations. For UP1, the interactions of the tandem RRM domains were mapped out (**Fig. S9 A,B**). The simulations revealed an interaction surface centered around the salt bridge interactions, but also encompassing the flexible N-terminus of RRM1, α_1_ on RRM2, and the loop between α_2_ and β_4_ on RRM2. These results present the possibility of RRM2 positioning the N-terminus and its 3_10_ helix for formation of the RNA binding pocket it forms with the inter-RRM linker. Identical simulations for UP1^dm^ revealed almost all inter-RRM interactions are ablated (**Fig. S9 C,D**). With the loss of the salt bridge interaction, RRM2 loses its interactions with the RRM1 N-terminus and can’t position it for formation of the RNA binding pocket with the inter-RRM linker, confirming previous simulations.

For simulations with full length hnRNP A1, the protein was divided into four domains: RRM1 (1-90), inter-RRM linker (91-106), RRM2 (107-182), and LCD_A1_ (183-320). The interactions for LCD_A1_ with each other domain was then analyzed. The simulations revealed LCD_A1_ shares a narrow interaction surface with the inter-RRM interface formed between the N-terminus of RRM1 and the α_2_/β_4_ loop of RRM2 (**Fig. S9 E,F,G**). Further contact with the tandem RRM domains were found with the β_2_/β_3_ loop of RRM1 and β_4_ of RRM2. With A1^dm^, the binding surface between RRM1 and LCD_A1_ was left primarily intact, with the exception of the RRM1 N-terminus (**Fig. S9 I,J**). For RRM2 (**Fig. S9K**), the loss of the salt bridge ablated all interactions between RRM2 and LCD_A1_, only a single interaction between K183 and E135 was detected. This result demonstrates that the allosteric coupling that exists between the tandem RRMs on hnRNP A1 propagates to its C-terminus domain.

For the inter-RRM linker, the interactions with LCD_A1_ were largely identical between hnRNP A1 (**Fig. S9H**) and A1^dm^ (**Fig. S9L**), with one significant difference. In hnRNP A1, the linker interacts with the RGG box region of LCD_A1_, whereas in A1^dm^, the RGG box residues shift from interacting with the linker to interacting with RRM1. They interact with two different regions of RRM1, residues 24-27, in the α-helix between β_1_ and β_2_, and residues 47-55, in the loop between β_2_ and β_3_.

### Thermodynamic coupling of the tandem RRMs of hnRNP A1 regulates its RNA binding capacity

To determine how the observed thermodynamic coupling of the tandem RRMs of hnRNP A1 impact RNA binding in cells, we next conducted PAR-CLIP studies for hnRNP A1 and the A1^dm^ variant. HEK293T cells were transiently transfected with myc-tagged hnRNP A1 and A1^dm^ constructs, treated with 4-thiouridine, UV-crosslinked and protein-RNA complexes were immunoprecipitated. Both hnRNP A1 and A1^dm^ were expressed well in cells and immunoprecipitated efficiently (**Fig. 4A**). Surprisingly, for hnRNP A1 we observed the presence of an additional higher molecular weight (HMW) protein band, likely corresponding to a dimer of ∼70kD, that immunoprecipitated in complex with higher levels of RNA compared with monomeric hnRNP A1 or the A1^dm^ (**Fig. 4A**). In two independent PAR-CLIP experiments, bound RNA was further purified away from hnRNP A1, hnRNP A1-HMW as well as monomeric A1^dm^ and sequenced. Reassuringly, PAR-CLIP-derived reads in all six libraries predominantly contained T-to-C substitutions, which is a result of mutations induced at the protein-RNA crosslinking site (**Fig. S10A**). PCA analysis of PAR-CLIP libraries revealed that the identities of bound RNA sequences for A1^dm^ separated from hnRNP A1 and hnRNP A1-HMW, which clustered together (**Fig. 4B**). Although consensus motif sequences identified by PAR-CLIP for hnRNP A1, hnRNP A1-HMW and the A1^dm^ variant were similar and AG-rich (**Fig. 4C**), comparative analysis of binding site occupancies across all transcripts pull-down show distinct variations. For example, hnRNP A1 and the HMW variant showed enriched binding to introns relative to the A1^dm^ mutant (**Fig. 4D, E, S10B**), whereas the A1^dm^ mutant more frequently associated with mRNAs and other RNA species (i.e. ncRNAs, snRNAs, miscRNAs etc.). Altogether, the difference in binding capacity between WT hnRNP A1 and the A1^dm^ mutant correlated with the latter being defective in its ability to dimerize on the immunoprecipitated transcripts.

**Figure 4.**
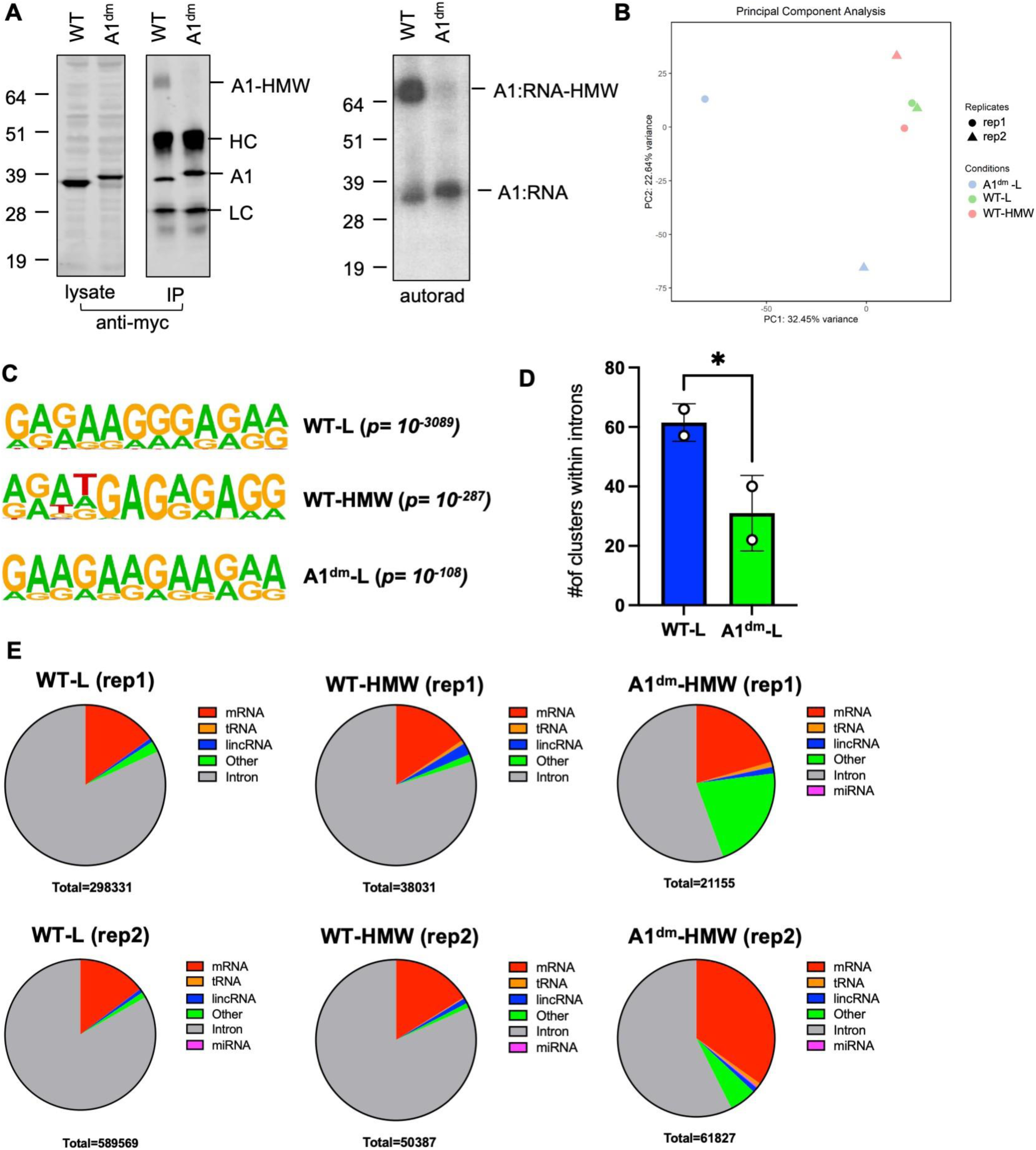
**PAR-CLIP analysis reveals distinct RNA-binding profiles of WT hnRNP A1 and the A1^dm^ mutant.** HEK293T cells transfected with myc-tagged WT hnRNP A1 and the A1^dm^ mutant were processed for PAR-CLIP. (A) Cell lysates and immunoprecipitated protein-RNA complexes were analyzed by immunoblotting using an anti hnRNP A1 antibody or autoradiography. HC and LC denote the heavy and light chain of the immunoprecipitating antibody. (B) PCA analysis of the 6 independent CLIP libraries. (C) Motif analysis of the RNA sequences bound by monomeric hnRNP A1 (L), high molecular weight (HMW) WT variant and the A1^dm^ mutant. (D) Number of intronic clusters amongst the top 100 most frequently bound clusters derived from PAR-CLIP are shown for WT hnRNP A1-L and the A1^dm^ mutant. (E) Number of PAR-CLIP reads that map to the indicated RNA species from the two independent PAR-CLIP libraries are shown.

## Discussion

### HnRNP A1 is an allosterically regulated RNA binding protein

Knowledge as to how hnRNP A1 converts recognition of short degenerate RNA sequence motifs, ubiquitous across a transcriptome, into regulated biological outcomes remains enigmatic. Numerous atomic-resolution structures and high-throughput binding studies offer insights into the mechanisms by which hnRNP A1 achieves specificity for a minimal 5’-YAG-3’ motif [1, 22]; however, these cumulative observations provide only limited understanding into the processes by which hnRNP A1 modulates general RNA metabolism. This study provides evidence that hnRNP A1 is in part intrinsically regulated via the thermodynamic coupling of its tandem RRM domains. The significance of intrinsic regulation is that hnRNP A1’s RNA binding capacity can be tuned by long-range communication between its tandem RRMS and its LCD, naturally occurring mutations, and post-translational modifications [2]. Indeed, more than 30 cancer associated mutations (∼15% of UP1 sequence) map to the tandem RRMs of hnRNP A1 with several having potential to disrupt the thermodynamic coupling described here (**Fig. 5**).

**Figure 5:**
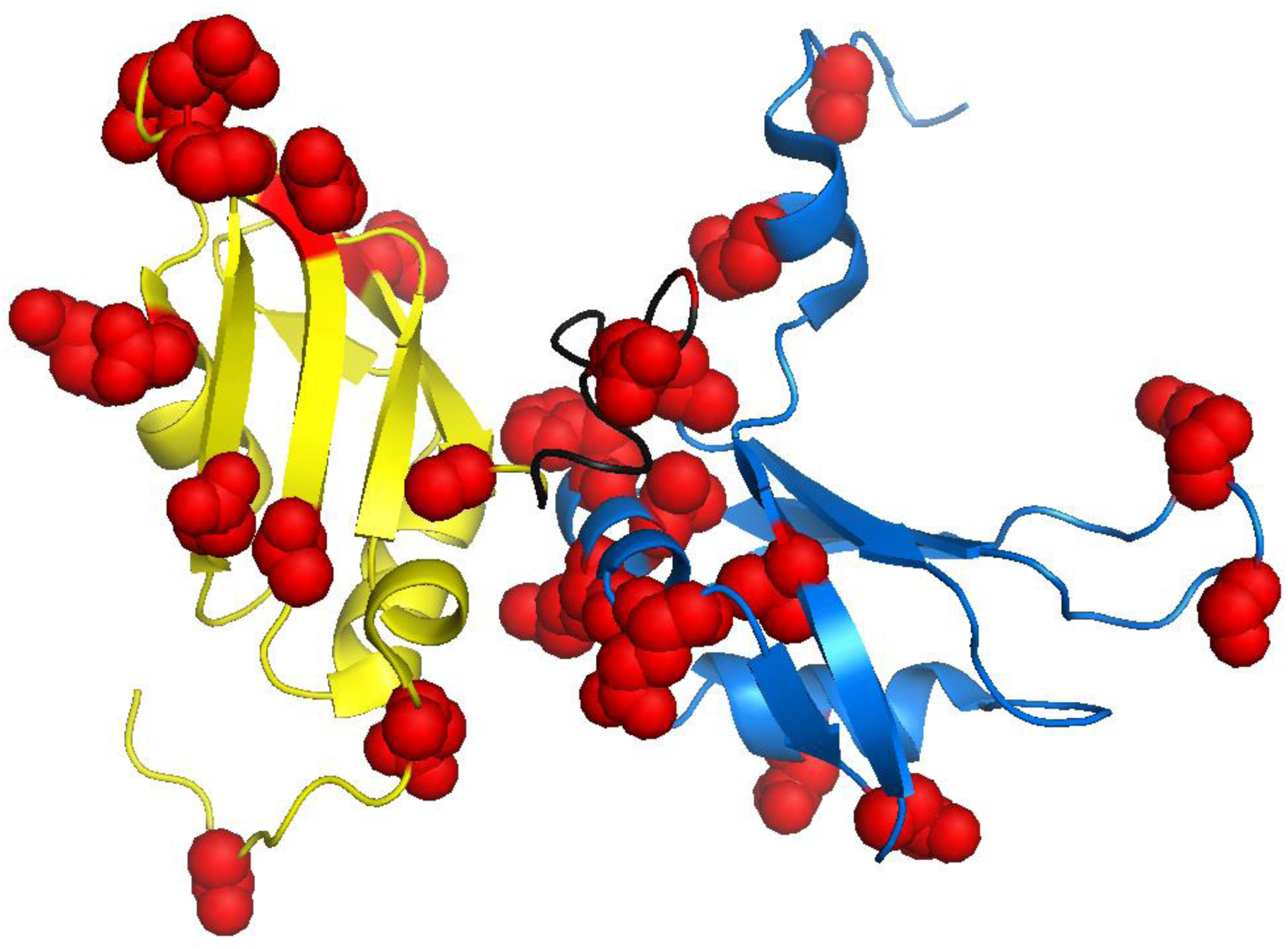
Cancer causing mutations in UP1 were compiled from the Cancer Cell Line Encyclopedia (D27N, D27E, E28K, P49S, N50D, R53H, G56S, T61A, V84L, P98L, V109I, D123N, H156R, V159L, K161E, Q165L, E185Q, S191C) and The Cancer Genome Atlas (S4L, E9R, D42N, R55M, A89V, Q96R, R140P, S142I, K161E, N171D, V177A. Residues identified in this compilation are shown as spheres and colored red. PDB:2lyv used for modeling.

In this study, we demonstrate that thermodynamic coupling of the tandem RRMs of hnRNP A1 regulates its RNA binding capacity and likely its ability to dimerize. Since specific RNA binding and dimerization are partitioned to the N-terminal UP1 domain and the LCD_A1_, respectively, we suggest that thermodynamic coupling of the tandem RRMs is the foundational basis of allosteric regulation of hnRNP A1. Based on the previously determined UP1-(AGU) crystal structure, we built a data-driven structural model of UP1 bound to HIV-1 SL3^ESS3^, a 25- nucleotide stem loop with a high affinity UAG binding site in its apical heptaloop [7]. In the model, the center of the β-sheet surface of RRM2 is more than 20 Å away from the heptaloop surface. The observation of this univalent binding mechanism opened the possibility that hnRNP A1 is an allosterically regulated RNA binding protein. Of further support of this concept, we demonstrated that mutating the R75:D155/R88:D157 salt bridge interactions, which stabilize the inter-RRM interface, were sufficient to reduce the affinity for SL3^ESS3^ by more than 18-fold with an accompanied decrease in total binding enthalpy [7]. While the large reduction in binding affinity alone does not satisfy conditions for allostery, the comparative high pressure NMR studies of wild type UP1 and the UP1^dm^ determined here clearly show that the integrity of the inter-RRM interface is necessary to impart uniform thermodynamic stability across the surface of the N-terminal domain of hnRNP A1. Decoupling the interface either via salt bridge mutations or RRM transposition exposes the significantly less stable RRM2 domain. Our results further revealed that the differential thermodynamic stabilities and conformational dynamics of RRM1 and RRM2 correlate with the extent of hydrophobic packing. Thus, this work demonstrates that the physicochemical basis of hnRNP A1 allostery is thermodynamic coupling, which acts to normalize the stabilities of the RRMs and to provide an intermolecular network to communicate binding site occupancy.

Allosteric regulation of hnRNP A1 offers a mechanistic rational to interpret its pleiotropic RNA binding functions. It is established that the LCD participates in heterotypic protein-protein and protein-RNA interactions, where the extent to which such interactions form influences the phase separation properties of hnRNP A1. Allosteric coupling of the tandem RRMs and the LCD_A1_ therefore allows fine-tuning of hnRNP A1’s heterotypic interactions via its specific affinity for RNA substrates that contain optimal or near optimal consensus motifs [22].

Related to this concept, the Mittag group found that the LCD_A1_ lays over the tandem RRMs [23]. This interaction is electrostatically driven, and dependent on the salt concentration of the solution buffer. This interaction drove stress granule assembly, as a lower salt buffer concentration led to greater inter-domain interaction and greater granule assembly. More recently, the Jeschke group performed PRE experiments with labeled residues in the LCD_A1_ to determine the points of contact between the two domains [24]. One such probe they used was located at residue 231 in the RGG box region. This probe showed clusters in elements important for RNA recognition such as the inter-RRM linker and the β2-β3 loop of RRM1. Since the LCD_A1_ domain occupies the same space as the RNA, it was hypothesized the RNA would displace LCD_A1_ from its binding surface. Our molecular dynamic simulations revealed similar results, as LCD_A1_ was found to interact with the inter-RRM linker and the RRM1 N-terminus, two elements that act in tandem to form a part of the RNA binding pocket. Our simulations also revealed the RGG box residues interact with the linker in WT hnRNP A1 but switch to interacting with the β_2_-β_3_ loop in the A1^dm^ mutant.

This previous research, along with our simulation and PAR-CLIP data, reveals that hnRNP A1 acts as a single functional unit through allosteric regulation. While each domain has its particular function, changes in one domain can be transmitted to others to affect their function. This can be seen in analysis of the post-translational modifications (PTMs) of hnRNP A1 (Table 1). Almost all PTMS occur in either RRM1 or LCD_A1_ and affect the function of the domain; RNA-binding for RRM1 and complex formation for LCD_A1_. A few PTMS affect the function of other domains. Confirmation of the regulatory role of RRM2 in RNA binding was found with ubiquitination of K183 [25]. The modification at this residue was found to disrupt the RNA-protein interaction by conformational changes in RRM2. In LCD_A1_ methylation of Arg218 and Arg225 decreases the ability of hnRNP A1 to activate IRES-mediated translation while also reducing its ITAF activity [26-28]. This regulation of identity-specific RNA binding activity could occur through the disruption of functionally specific complex formation required to direct hnRNP A1 to its RNA substrate. This RNA complex-dependent regulation could explain our observation through PAR-CLIP that the A1^dm^ mutant, which loses inter-domain coupling, binds RNA of different identity.

**Table 1:** Compilation of post-translational modifications made to hnRNP A1 and their locations on the protein. (Adapted from Clarke et al)

| Modification | RRM1 | RRM2 | LCD |
| --- | --- | --- | --- |
| Phosphorylation | Serine 4[30] | Serine 192[31] | Serine 199[32] |
|  | Serine 6[30] |  | Serine 310-312[31] |
|  | Threonine 51[29] |  |  |
|  | Serine 95[33] |  |  |
| Methylation | Arginine 31[34] |  | Arginine 206[28] |
|  |  |  | Arginine 218[26] |
|  |  |  | Arginine 225[27] |
|  |  |  | Arginine 232[28] |
| Ubiquitination | Lysine 3[35] | Lysine 183[25] | Lysine 298[25] |
|  | Lysine 8[35] |  |  |
|  | Lysine 15[35] |  |  |
| Acetylation | Lysine 3[36] |  |  |
|  | Lysine 52[36] |  |  |
|  | Lysine 87[36] |  |  |
| SUMO-ylation |  | Lysine 183[30] |  |
| β-N-acetyl-glucosamine-ylation | Serine 22[37] | | |
| PAR-ylation |  |  | Lysine 298[38] |

One of the major questions presented by our findings is what specific advantage does hnRNP A1 gain from having a stable RRM1 appended to an unstable RRM2. The stability of hnRNP A1 is important physiologically, as the protein’s stability plays a role in cellular survival. It has been shown that as a response to endoplasmic reticulum stress that eukaryotic initiation translation factor 3 alpha kinase 3 (PERK) phosphorylates Thr51 on RRM1, leading to its degradation [29]. This degradation occurs with its unfolding, potentially aided by its coupling to the unstable RRM2 domain.

When combining these various observations, it becomes more likely that allostery intrinsically underlies the regulatory capacity of hnRNP A1 to engage with its various RNA targets. Allostery is a well-established property of multi-domain enzymes, which coordinate multiple biomolecular inputs to catalyze functional outcomes [17]. Indeed, enzymes that function in primary metabolism are almost universally allosterically controlled by substrate binding and post-translational modifications. Efforts to identify and characterize potential allosteric control of RNA binding proteins that recognize short degenerate sequences might offer critical insights into RNA biology. Further, this work demonstrates how allostery can be masked via thermodynamic coupling of domains and the importance of employing various biophysical methods to elucidate it.

## Materials and Methods

### Constructs and protein purification

All protein constructs were cloned from gBlock gene fragments (IDT) into pMCSG plasmid [39]. UP1 and RRM constructs were cloned with an N-terminal 22 amino acid His-Tag while full length hnRNP A1 constructs were cloned with an N-terminal GB1 solubility domain with a His-Tag. All constructs contained TEV cleavage sites for removal of the His-Tag and GB1 domain.

Protein was overexpressed as previously described [21]. UP1 and RRM proteins were purified as previously described [40, 41]. Full length hnRNP A1 was purified in similar manner, but with lysis buffer composition of 20 mM Na_2_HPO_4_ pH=7.5, 1.2 M NaCl, 0.5 mM EDTA, 2 M Urea, 20 mM imidazole, 1 mM PMSF. Elution buffer composition was identical except imidazole concentration was 250 mM. FPLC Gel Filtrations buffer was 100 mM HEPES pH=7.5 and 1 M NaCl. Salt concentration was decreased by serial dilutions with 100 mM HEPES pH=7.5 while simultaneously concentrating protein sample through Amicon filtration.

### Go-model simulations

List of native contacts for the all-atom simulations with a structure-based potential (Go-model) was established with SMOG algorithm [42] using the coordinates of RRM1 and RRM2 extracted from PDB 1U1R as a reference structure. All-atom simulations were performed in GROMACS 2018.8 using the leapfrog integration method with 2 fs timesteps [43]. For each RRM, independent simulations were performed over a wide range of temperatures and the folding temperature (T_f_) was determined with the weighted histogram analysis method (WHAM) [44]. Analysis of the trajectories was carried out using in-house python scripts.

### High-pressure NMR

All NMR spectra were acquired on a Bruker 700 spectrometer equipped with z-shielded gradient triple resonance 5 mm TCI cryoprobe. Hydrostatic pressure was controlled using a commercial ceramic high-pressure NMR cell and an automatic pump system (Daedalus Innovations, Philadelphia, PA). 2D ^1^H-^15^N HSQC experiments were recorded at 290K with a time domain matrix consisting of 100* (t_1_, ^15^N) × 1024* (t_2_, ^1^H) complex points with acquisition time of 50 ms (t_1_) and 91.8 ms (t_2_) using 1.5 s interscan delay. 2D ^1^H-^15^N spectra were collected every 250 bar from 1 bar to 2.5 kbar using 20 min equilibration delay after every change of pressure. All spectra were processed using NMRPipe [45] and displayed with SPARKY [46].

### Relaxation Dispersion

NMR relaxation dispersion experiments were acquired on a 600 MHz Bruker magnet equipped with a 5 mm TXI cryoprobe at 298 K. Amide ^15^N CPMG experiments were acquired at using the STCW-CPMG [47] pulse sequences and recorded in a pseudo-3D fashion. The constant relaxation time was set to 40 ms and the CPMG pulsing frequency, *ν*_CPMG_, was varied from 25 Hz to 1.5 kHz. To prevent spectral artifacts arising from sample instability, the order of the acquisition of increments along the indirect t1 dimension was randomized and interleaved with the number of scans, repeating the minimal phase cycle four times. NMR data was processed using nmrPipe [45] and intensities were extracted using the autoFit routine in the NMRPipe suite. Errors in signal amplitudes were estimated from 2-3 replicate *ν*_CPMG_ measurements for each CPMG experiment and used to propagate the errors in *R*_2,eff_, where

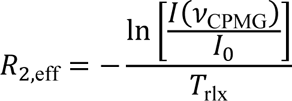

*I*_0_ is the signal amplitude without the CPMG element, *I*(*ν*_CPMG_) is the amplitude at the given *ν*_CPMG_, and *T*_rlx_ is the constant relaxation time. The median error in *R*_2,eff_ was 0.4, 1.6, and 3.3 s^-1^ for ^15^N datasets of RRM1, RRM2, and UP1, respectively. The ^15^N CPMG profiles were analyzed to identify those residues with *R*_ex_ = *R*_2,eff_(min(*ν*_CPMG_)) – *R*_2,eff_(max(*ν*_CPMG_)) greater than 1.65 times the *R*_ex_ measurement error, σ*_R_*_ex_, amounting to 95% confidence for the presence of exchange. Those residues identified in this fashion were then fit to simple analytical models of fast and slow exchange [48] to estimate the initial 2-state exchange parameters, the total rate *k*_ex_ = *k*_ab_ + *k*_ba_ and population *p*_b_. All exchange was in the fast regime with *k*_ex_ on the order of ∼200 – 2000/s with the most accurate results for RRM1 of about 175/s. Populations of the excited state could not be obtained. Processed spectra were displayed with SPARKY [46].

### High-pressure SAXS

The HP-SAXS data were collected at the Sector 7A1 station (HP-Bio) of the Cornell High Energy Synchrotron Source (CHESS) [49-51]. Pressure was maintained by a Barocycler HUB440 high-pressure pump (Pressure BioSciences). Samples were illuminated with a 250 × 250-µm X-ray beam of wavelength λ = 0.8823 Å (14.05 keV), with a flux of ∼2.1 × 10^11^ photons/s. Scattering was measured with an EIGER 4M detector (Dectris) in vacuum with a pixel size of 75 × 75 µm and an active area of 155.1 × 162.2 mm (2068 × 2162 pixels), over the *q* range of ∼0.01–0.6 Å^−1^ [where *q* is the wave vector defined as *q* = (4π sin*θ*/*λ*) and 2*θ* is the scattering angle].

Samples were prepared in 100 mM HEPES pH 7.5, 350 mM NaCl, 0.5 mM EDTA at 5 mg/mL for A1^dm^ and 10 mg/ml for wt hnRNP A1. HP-SAXS measurements were carried on from 1 bar to 2.5 kbar at 290 K with 10 exposures of 1.0 s each (for a total of 10-s exposure at each dataset). The 2D SAXS images were azimuthally integrated about the beam center and normalized by the transmitted intensities via standard image correction procedures using the BioXTAS RAW 2.1.4 software package [52]. Data was placed on an absolute scale using water as a standard. The specimen-to-detector distance was set to 1804.0 mm. Data analysis was performed with BioXTAS RAW 2.1.4 [52].

### PAR-CLIP

HEK 293T cells were grown in Dulbecco’s Modified Eagle Medium (DMEM) supplemented with 10% fetal bovine serum. HEK293T were seeded in 10-cm dishes then transfected with 10 ug of myc-tagged hnRNPA1 and hnRNP A1^dm^ mutant using polyethyleneimine (PolySciences). Cells were treated with 100 uM 4-SU 16 hours prior to UV-crosslinking. PAR-CLIP was performed as previously described [53]. In brief, cells were lysed in 1x RIPA buffer supplemented with protease inhibitors. hnRNP A1-RNA complexes were immunoprecipitated by a rabbit polyclonal antibody against the myc tag (Thermo-Fisher, #PA1-981) and the bound RNA molecules end-labeled with T4 PNK and γ-^32^P-ATP. Protein-RNA complexes were resolved on 4- 12% NuPAGE SDS-PAGE gels, transferred to nitrocellulose membranes and visualized by autoradiography. Bound RNA was further purified away from protein-RNA complexes by proteinase K treatment and processed by sequential ligation of 3’ and 5’ adapters. Resulting libraries were subjected to reverse transcription, PCR and sequenced on an Illumina Next-seq instrument for 75 cycles.

### Analysis of PAR-CLIP data

PAR-CLIP derived reads were first processed using BBduk to remove adapter sequences and separated into barcodes, removing reads of length less than ten nucleotides. The resulting reads were mapped to rRNA, then the unmapped reads were mapped to the human transcriptome using STAR aligner, allowing mismatch in less than 10% of read length. PCA analysis was performed as described before [54]. Mapped reads were further analyzed by R package wavClusteR for the presence of T-to-C substitutions and by PARalyzer to generate clusters (i.e. binding sites) and annotated with annotatePeaks.pl by HOMER.

### Molecular Dynamic Simulations

The full length hnRNP A1 protein (1-320) was developed using AlphaFold [20]. Coordinates for hnRNPA1 R75D/R88D mutant PDB were designed via pymol and based on the initial orientation of full-length hnrnp A1 (1-320). The protonation of the protein and the fine-tuning of the hydrogen-bond network were facilitated through the Protein Prepare module in HTMD [55].

The systems were prepared for simulation via HTMD, with water padding configured at a distance equivalent to the furthest atom of the protein plus an extra 10 Å from the center. The system was neutralized with the addition of sodium and chloride ions. For systems, an equilibration process was implemented, followed by production cycles of 1 μs each per system.

The equilibration protocol comprised 500 steps each of minimization and NVT, followed by 1 ns NPT. Heavy atoms were subjected to restraints of 1 kcal/mol, and nonheavy atoms had restraints of 0.1 kcal/mol, which were progressively lifted until midway through the simulation when the system became entirely free of restraints.

The production simulations ran without restraints for a duration of 1us. The CHARMM22* force field was employed, with simulations executed on a cluster equipped with a single GPU, utilizing ACEMD simulation software on NMRBOX [56]. Using Bridg2, an analysis was conducted on the hydrogen-bond network and its occupancy over the 1us trajectory [57]. The objective was to uncover any potential interactions between the RRM domains, inter-RRM linker, and C-terminal domain within the established hydrogen-bond network of WT hnRNP A1 and R75D/R88D mutant. Through the Bridg2’s built in tools, the hydrogen-bond network graph and its occupancy were established based on the residue numbers pertaining to the specific domains.

## Acknowledgements

The high-pressure SAXS experiments were conducted at the Center for High-Energy X-ray Sciences (CHEXS), which is supported by the National Science Foundation (BIO, ENG and MPS Directorates) under award DMR-1829070, and the Macromolecular Diffraction at CHESS (MacCHESS) facility, supported by award 1-P30-GM124166-01A1 from the National Institute of General Medical Sciences, National Institutes of Health, and by New York State’s Empire State Development Corporation (NYSTAR). We would like to thank Dr. Richard Gillilan and Dr. Qingqiu Huang for their technical support. The CLIP-seq data has been deposited to the GEO server (accession number: GSE238202). This work was additionally funded by the Center for HIV RNA studies, U54 AI170660 (SBK and BST) and R01AI150830 (BST).

## Author Contributions

Conceptualization (BST and JDL); Material preparation (JDL): HP NMR data collection and analysis (JR): Go-model simulations (DP): NMR dynamics data collection and analysis (JDL, JR, and ALH); Molecular dynamics simulations (SP); and PAR-CLIP analysis (SK, AK and YW); Writing and editing manuscript (BST, JDL, JR and SK)

## References

1. Levengood, J.D. and B.S. Tolbert, Idiosyncrasies of hnRNP A1-RNA recognition: Can binding mode influence function. Semin Cell Dev Biol, 2019. 86: p. 150–161.

2. Clarke, J.P., et al., A Comprehensive Analysis of the Role of hnRNP A1 Function and Dysfunction in the Pathogenesis of Neurodegenerative Disease. Front Mol Biosci, 2021. 8: p. 659610.

3. Jean-Philippe, J., S. Paz, and M. Caputi, hnRNP A1: the Swiss army knife of gene expression. Int J Mol Sci, 2013. 14(9): p. 18999–9024.

4. Geuens, T., D. Bouhy, and V. Timmerman, The hnRNP family: insights into their role in health and disease. Hum Genet, 2016. 135(8): p. 851–67.

5. Han, S.P., Y.H. Tang, and R. Smith, Functional diversity of the hnRNPs: past, present and perspectives. Biochem J, 2010. 430(3): p. 379–92.

6. Cartegni, L., Maconi, M., Morandi, E., Cobianchi, F., Riva, S., and Biamonti, G., hnRNP A1 selectively interacts through its Gly-rich domain with different RNA-binding proteins. J Mol Biol, 1996. 259: p. 337–348.

7. Morgan, C.E., et al., The First Crystal Structure of the UP1 Domain of hnRNP A1 Bound to RNA Reveals a New Look for an Old RNA Binding Protein. J Mol Biol, 2015. 427(20): p. 3241–3257.

8. Ding, J., Hayashi, M.K., Zhang, Y., Manche, L., Krainer, A.R., and Xu, R.M., Crystal structure of the two RRM-domain of hnRNP A1 (UP1) complexed with single-stranded telomeric DNA. Genes Dev., 1999. 13: p. 1102–1115.

9. Beusch, I., et al., Tandem hnRNP A1 RNA recognition motifs act in concert to repress the splicing of survival motor neuron exon 7. Elife, 2017. 6.

10. Kooshapur, H., et al., Structural basis for terminal loop recognition and stimulation of pri-miRNA-18a processing by hnRNP A1. Nat Commun, 2018. 9(1): p. 2479.

11. Vitali, J., Ding, J., Jiang, J., Zhang, Y., Krainer, A.R., and Xu, R.M., Correlated alternative side chain conformations in the RNA-recognition motif of heterogeneous nuclear ribonucleoprotein A1. Nucleic Acids Res, 2002. 30(7): p. 1531–1538.

12. Barraud, P. and F.H. Allain, Solution structure of the two RNA recognition motifs of hnRNP A1 using segmental isotope labeling: how the relative orientation between RRMs influences the nucleic acid binding topology. J Biomol NMR, 2013. 55(1): p. 119–38.

13. Myers, J.C. and Y. Shamoo, Human UP1 as a model for understanding purine recognition in the family of proteins containing the RNA recognition motif (RRM). J Mol Biol, 2004. 342(3): p. 743–56.

14. Noel, J.K., Onuchic, J.N., The many faces of structure-based potentials: from protein folding landscapes to structural characterization of complex biomolecules. Dokholyan NV (ed) Computational modeling of biological systems. 2012: Springer, New York, NY.

15. Hills, R.D., Jr. and C.L. Brooks, 3rd, Insights from coarse-grained Go models for protein folding and dynamics. Int J Mol Sci, 2009. 10(3): p. 889–905.

16. Singh, A., J.A. Purslow, and V. Venditti, 15N CPMG Relaxation Dispersion for the Investigation of Protein Conformational Dynamics on the micros-ms Timescale. J Vis Exp, 2021(170).

17. Farber, P.J. and A. Mittermaier, Relaxation dispersion NMR spectroscopy for the study of protein allostery. Biophys Rev, 2015. 7(2): p. 191–200.

18. Kitahara, R., et al., Pressure-induced chemical shifts as probes for conformational fluctuations in proteins. Prog Nucl Magn Reson Spectrosc, 2013. 71: p. 35–58.

19. Roche, J., C.A. Royer, and C. Roumestand, Monitoring protein folding through high pressure NMR spectroscopy. Prog Nucl Magn Reson Spectrosc, 2017. 102-103: p. 15–31.

20. Jumper, J., et al., Highly accurate protein structure prediction with AlphaFold. Nature, 2021. 596(7873): p. 583–589.

21. Levengood, J.D., et al., Thermodynamic stability of hnRNP A1 low complexity domain revealed by high-pressure NMR. Proteins, 2021. 89(7): p. 781–791.

22. Jain, N., et al., Rules of RNA specificity of hnRNP A1 revealed by global and quantitative analysis of its affinity distribution. Proc Natl Acad Sci U S A, 2017. 114(9): p. 2206–2211.

23. Martin, E.W., et al., Interplay of folded domains and the disordered low-complexity domain in mediating hnRNPA1 phase separation. Nucleic Acids Res, 2021. 49(5): p. 2931–2945.

24. Ritsch, I., et al., Phase Separation of Heterogeneous Nuclear Ribonucleoprotein A1 upon Specific RNA-Binding Observed by Magnetic Resonance. Angew Chem Int Ed Engl, 2022. 61(40): p. e202204311.

25. Wang, F., et al., SPSB1-mediated HnRNP A1 ubiquitylation regulates alternative splicing and cell migration in EGF signaling. Cell Res, 2017. 27(4): p. 540–558.

26. Gao, G., S. Dhar, and M.T. Bedford, PRMT5 regulates IRES-dependent translation via methylation of hnRNP A1. Nucleic Acids Res, 2017. 45(8): p. 4359–4369.

27. Barrera, A., et al., Post-translational modifications of hnRNP A1 differentially modulate retroviral IRES-mediated translation initiation. Nucleic Acids Res, 2020. 48(18): p. 10479–10499.

28. Wall, M.L. and S.M. Lewis, Methylarginines within the RGG-Motif Region of hnRNP A1 Affect Its IRES Trans-Acting Factor Activity and Are Required for hnRNP A1 Stress Granule Localization and Formation. J Mol Biol, 2017. 429(2): p. 295–307.

29. Koo, J.H., et al., Endoplasmic Reticulum Stress in Hepatic Stellate Cells Promotes Liver Fibrosis via PERK-Mediated Degradation of HNRNPA1 and Up-regulation of SMAD2. Gastroenterology, 2016. 150(1): p. 181–193 e8.

30. Roy, R., et al., hnRNPA1 couples nuclear export and translation of specific mRNAs downstream of FGF-2/S6K2 signalling. Nucleic Acids Res, 2014. 42(20): p. 12483–97.

31. Buxade, M., et al., The Mnks are novel components in the control of TNF alpha biosynthesis and phosphorylate and regulate hnRNP A1. Immunity, 2005. 23(2): p. 177–89.

32. Martin, J., et al., Phosphomimetic substitution of heterogeneous nuclear ribonucleoprotein A1 at serine 199 abolishes AKT-dependent internal ribosome entry site-transacting factor (ITAF) function via effects on strand annealing and results in mammalian target of rapamycin complex 1 (mTORC1) inhibitor sensitivity. J Biol Chem, 2011. 286(18): p. 16402–13.

33. Sui, J., et al., DNA-PKcs phosphorylates hnRNP-A1 to facilitate the RPA-to-POT1 switch and telomere capping after replication. Nucleic Acids Res, 2015. 43(12): p. 5971–83.

34. Hsu, M.C., et al., Protein Arginine Methyltransferase 3 Enhances Chemoresistance in Pancreatic Cancer by Methylating hnRNPA1 to Increase ABCG2 Expression. Cancers (Basel), 2018. 11(1).

35. Fang, J., et al., Ubiquitination of hnRNPA1 by TRAF6 links chronic innate immune signaling with myelodysplasia. Nat Immunol, 2017. 18(2): p. 236–245.

36. Yang, H., et al., Sirtuin-mediated deacetylation of hnRNP A1 suppresses glycolysis and growth in hepatocellular carcinoma. Oncogene, 2019. 38(25): p. 4915–4931.

37. Hart, G.W., et al., Cross talk between O-GlcNAcylation and phosphorylation: roles in signaling, transcription, and chronic disease. Annu Rev Biochem, 2011. 80: p. 825–58.

38. Duan, Y., et al., PARylation regulates stress granule dynamics, phase separation, and neurotoxicity of disease-related RNA-binding proteins. Cell Res, 2019. 29(3): p. 233–247.

39. Stols, L., et al., A new vector for high-throughput, ligation-independent cloning encoding a tobacco etch virus protease cleavage site. Protein Expr Purif, 2002. 25(1): p. 8–15.

40. Levengood, J.D., et al., Solution structure of the HIV-1 exon splicing silencer 3. J Mol Biol, 2012. 415(4): p. 680–98.

41. Rollins, C., et al., Thermodynamic and phylogenetic insights into hnRNP A1 recognition of the HIV-1 exon splicing silencer 3 element. Biochemistry, 2014. 53(13): p. 2172–84.

42. Noel, J.K., et al., SMOG 2: A Versatile Software Package for Generating Structure-Based Models. PLoS Comput Biol, 2016. 12(3): p. e1004794.

43. Abraham, M.J., et al., GROMACS: High performance molecular simulations through multi-level parallelism from laptops to supercomputers. SoftwareX, 2015. 1-2: p. 19–25.

44. Kumar, S., Bouzida, D., Swendsen, R.H., Kollman, P.A., and Rosenberg, J.M., The weighted histogram analysis method for free-energy calculations on biomolecules. I. The method. J. Comput. Chem., 1992. 13(8): p. 1011–1021.

45. Delaglio, F., Grzesiek, S., Vuister, G.W., Zhu, G., Pfeifer, J., and Bax, A., NMRPipe: a multidimensional spectral processing system based on UNIX pipes. J Biomol NMR, 1995. 6(3): p. 277–93.

46. Goddard, D., and Kneller, D.G., Sparky 3. University of California, San Francisco, 2000.

47. Jiang, B., et al., A (15)N CPMG relaxation dispersion experiment more resistant to resonance offset and pulse imperfection. J Magn Reson, 2015. 257: p. 1–7.

48. Palmer, A.G., 3rd, C.D. Kroenke, and J.P. Loria, *Nuclear magnetic resonance methods for quantifying microsecond-to-millisecond motions in biological macromolecules*. Methods Enzymol, 2001. 339: p. 204–38.

49. Acerbo, A.S., M.J. Cook, and R.E. Gillilan, Upgrade of MacCHESS facility for X-ray scattering of biological macromolecules in solution. J Synchrotron Radiat, 2015. 22(1): p. 180–6.

50. Ando, N., et al., High hydrostatic pressure small-angle X-ray scattering cell for protein solution studies featuring diamond windows and disposable sample cells. Journal of Applied Crystallography, 2008. 41(1): p. 167–175.

51. Gillilan, R.E., High-pressure SAXS, deep life, and extreme biophysics. Methods Enzymol, 2022. 677: p. 323–355.

52. Hopkins, J.B., R.E. Gillilan, and S. Skou, BioXTAS RAW: improvements to a free open-source program for small-angle X-ray scattering data reduction and analysis. J Appl Crystallogr, 2017. 50(Pt 5): p. 1545–1553.

53. Shema Mugisha, C., K. Tenneti, and S.B. Kutluay, Clip for studying protein-RNA interactions that regulate virus replication. Methods, 2020. 183: p. 84–92.

54. Puray-Chavez, M., Lee, N., Tenneti, K., Wang, Y., Vuong, H.R., Liu, Y., Horani, A., Huang, T., Gunsten, S.P., Case, J.B., Yang, W., Diamond, M.S., Brody, S.L., Dougherty, J., and Kutluay, S.B., The translational landscape of SARS-CoV-2-infected cells reveals suppression of innate immune genes. mBIO, 2022. 13(3).

55. Harvey, M.J., Giupponi, G., and De Fabritilis, ACEMD: Accelerating biomolecular simulations in the microsecond time scale. J. Chem. Theor. Comput., 2009. 5: p. 1632–1639.

56. Maciejewski, M.W., et al., NMRbox: A Resource for Biomolecular NMR Computation. Biophys J, 2017. 112(8): p. 1529–1534.

57. Siemers, M. and A.-N. Bondar, Interactive Interface for Graph-Based Analyses of Dynamic H-Bond Networks: Application to Spike Protein S. Journal of Chemical Information and Modeling, 2021. 61(6): p. 2998–3014.

