## Supplemental Information for "Thermodynamic Coupling of the tandem RRM domains of hnRNP A1 underlie its Pleiotropic RNA Binding Functions"

**B.** This work was funded by the Center for HIV RNA studies, U54 AI170660 (SBK and BST) and R01AI150830 (BST).

**C.** Jeffrey D. Levengood[1], Davit Potoyan[2], Srinivas Penumutchu[1], Abhishek Kumar[3], Yiqing Wang[3], Alexandar L. Hansen[4], Sebla Kutluay[3], Julien Roche[5*], and Blanton S. Tolbert[1*]

[1]Department of Biochemistry and Biophysics, University of Pennsylvania, Pennsylvania, PA 19104 United States
[2]Department of Chemistry, Iowa State University, Ames, IA 50011, United States.
[3]Department of Molecular Microbiology, Washington University School of Medicine, St. Louis, MO 63110, United States.
[4]CCIC and Gateway NMR Facility, The Ohio State University, Columbus, OH, 43210, United States
[5]Roy J. Carver Department of Biochemistry, Biophysics and Molecular Biology, Iowa State University, Ames, IA 50011, United States.

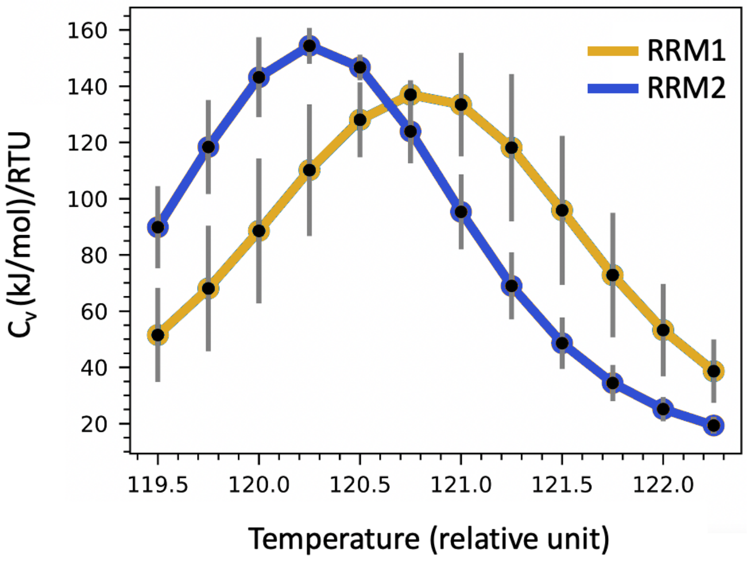


**Figure S1.** Specific heat capacity measured from all-atom molecular simulations conducted on the individual RRMs with a structure-based (Go-model) potential. These simulations predict that the isolated RRM1 (yellow) is thermodynamically more stable than the isolated RRM2 (blue).

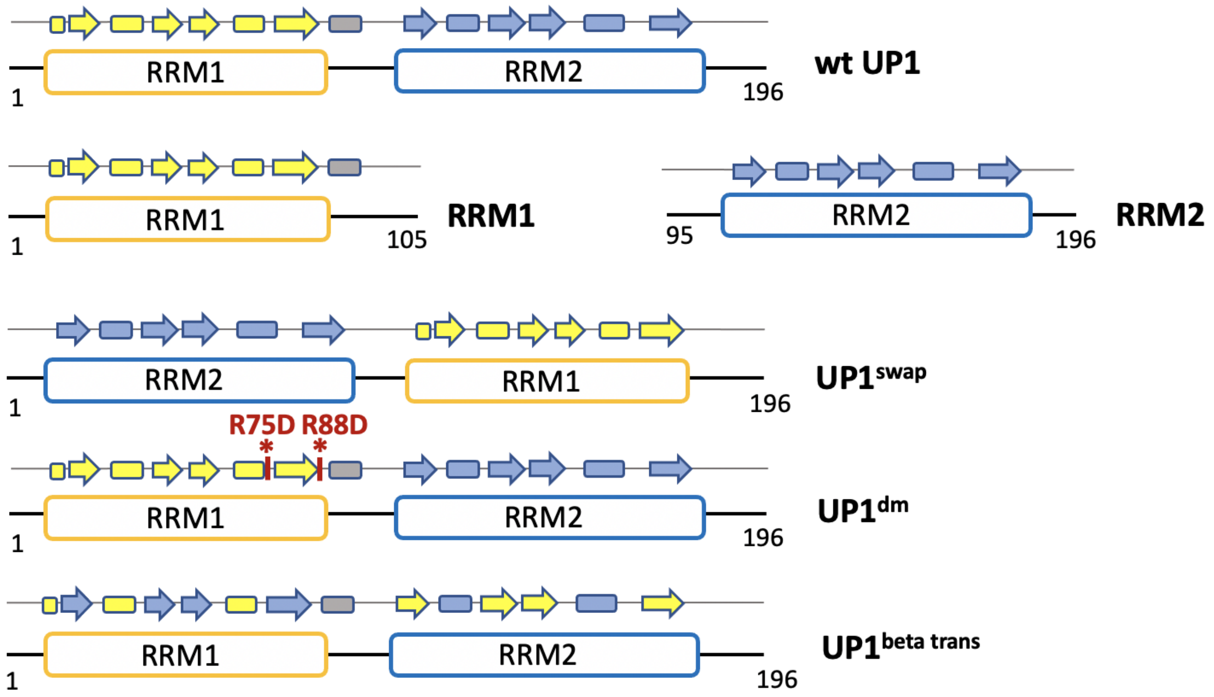


**Figure S2**. List of constructs used in this study, including from top to bottom: (*i*) wt UP1 domain of hnRNP A1, (*ii*) the isolated RRM1 and RRM2 motifs, (*iii*) UP1$^{swap}$ variant for which the RRM2 motif is positioned at the N-terminus and RRM1 at the C-terminus, (*iv*) UP1$^{dm,}$ a construct bearing two mutations (R75D, R88D) designed to disrupt two salt bridges at the interface between the two RRMs, and (*v*) UP1$^{beta\text{-}trans}$ for which the sequences of the four β-strands are swapped between RRM1 and RRM2.

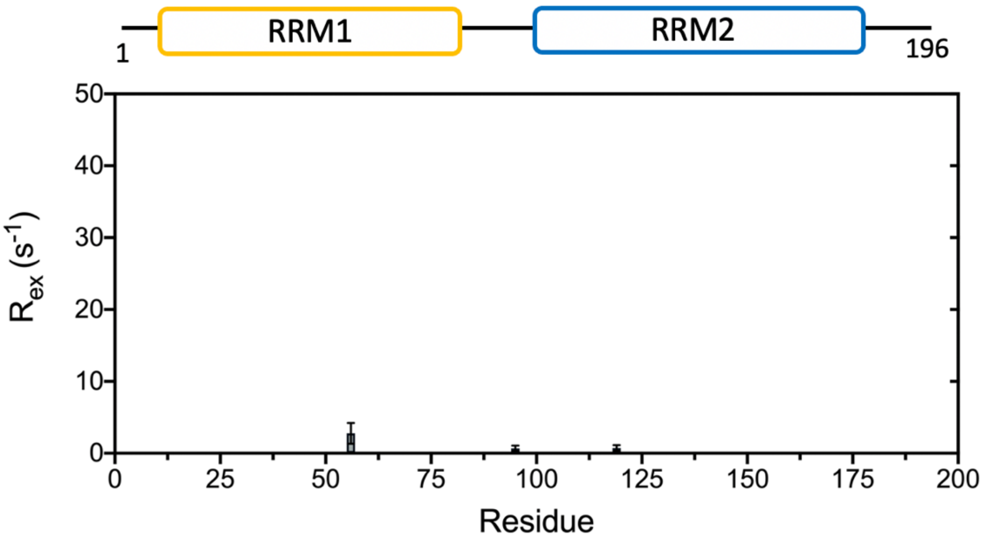


**Figure S3.** Chemical exchange contribution to the $^{15}N$ transverse relaxation rate ($R_{ex}$) measured for amide resonances of the isolated UP1 domain.

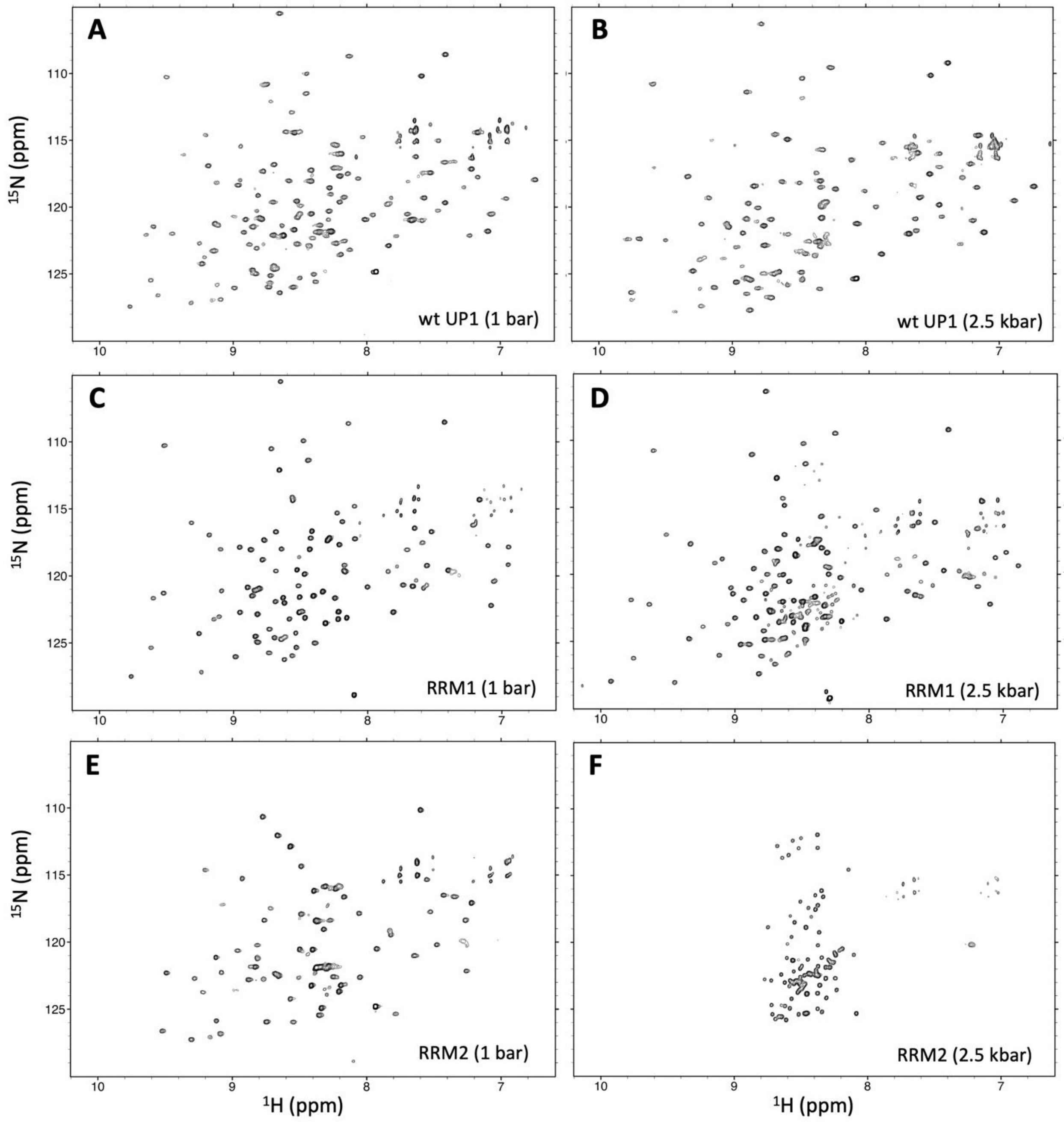


**Figure S4.** $^{15}N$-$^{1}H$ HSQC spectra collected at 1 bar and 2.5 kbar for (**A-B**) wt UP1, (**C-D**) the isolated RRM1 motif, and (**E-F**) the isolated RRM2 motif. Spectrum with narrow $^{1}H$ chemical shift dispersion observed for RRM2 at 2.5 kbar is characteristic of fully disordered protein chains and therefore indicates that RRM2 experiences complete unfolding within this pressure range.

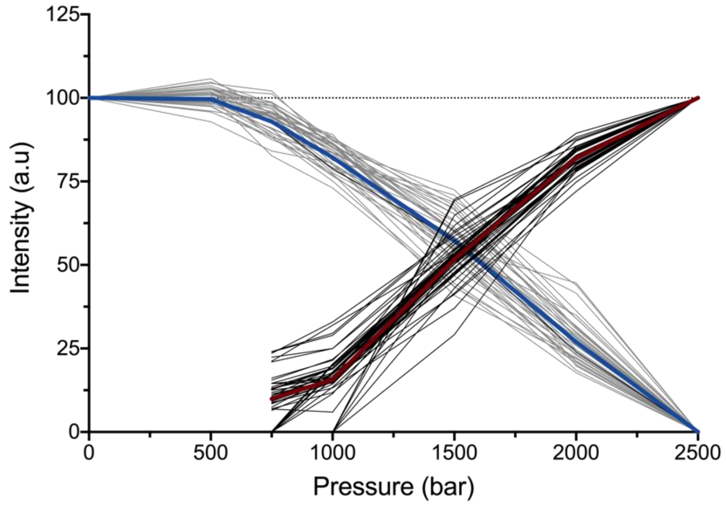


**Figure S5**. Comparison of peak intensity profiles measured for amide resonances assigned to residues of RRM2 in a native (folded) state (pale gray lines, average profile shown in blue) with intensity profiles measured residues of RRM2 in a non-native (unfolded) state (black lines, average profile shown in red).

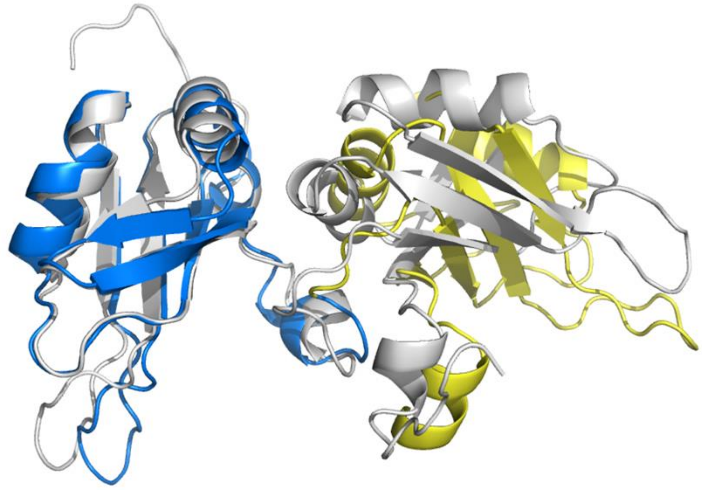

**Figure S6**. Comparison of the structure prediction generated by Alphafold2 for the UP1$^{swap}$ variant (with RRM2 in blue and RRM1 in yellow) with the reference high-resolution X-ray structure of UP1 (gray, pdb 1U1R).

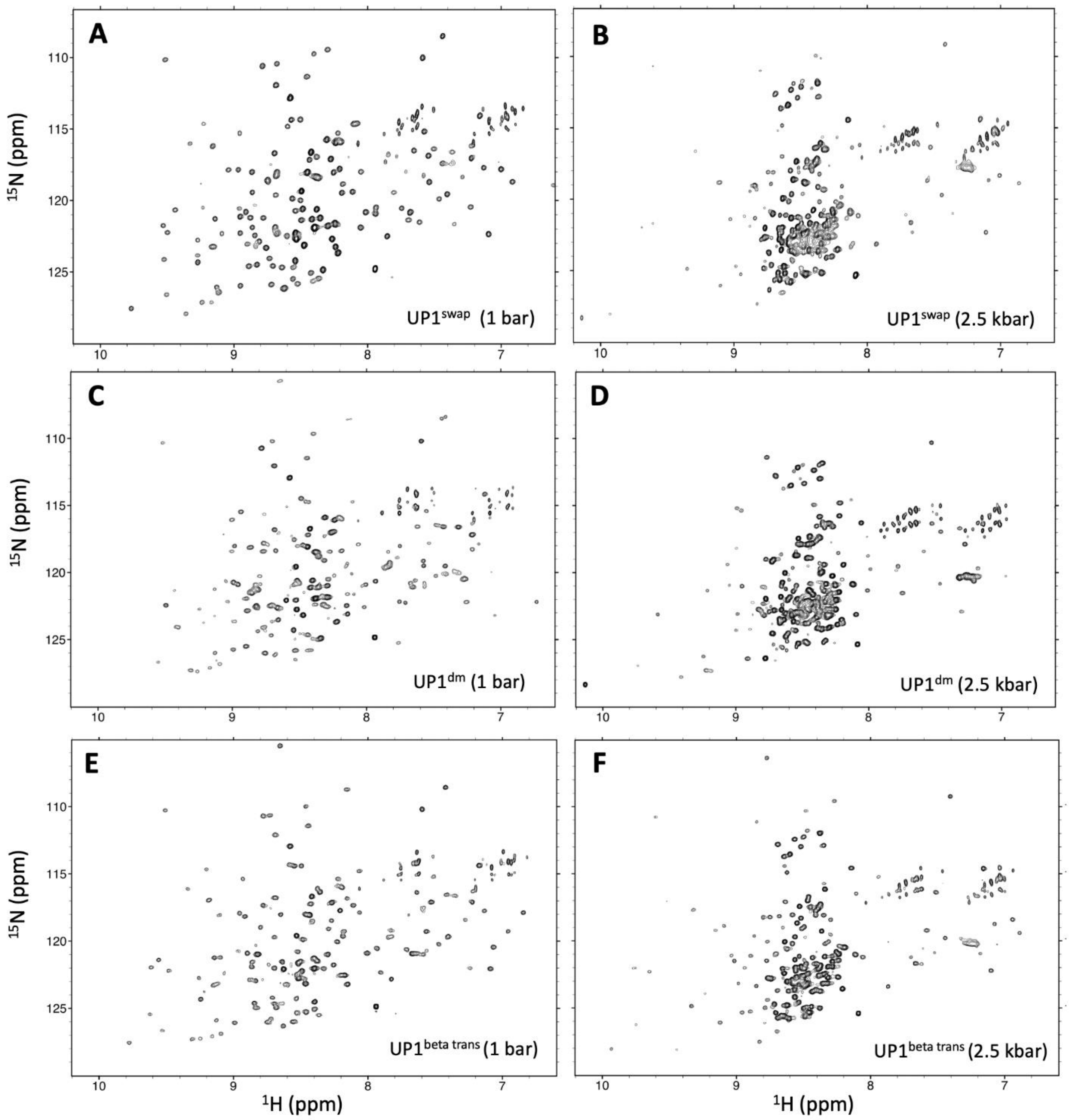


**Figure S7.** $^{15}$N-$^{1}$H HSQC spectra collected at 1 bar and 2.5 kbar for (**A-B**) UP1$^{swap}$, (**C-D**) UP1$^{dm}$, and (**E-F**) UP1$^{beta\ trans}$. All three variants show evidence of significant unfolding under pressure.

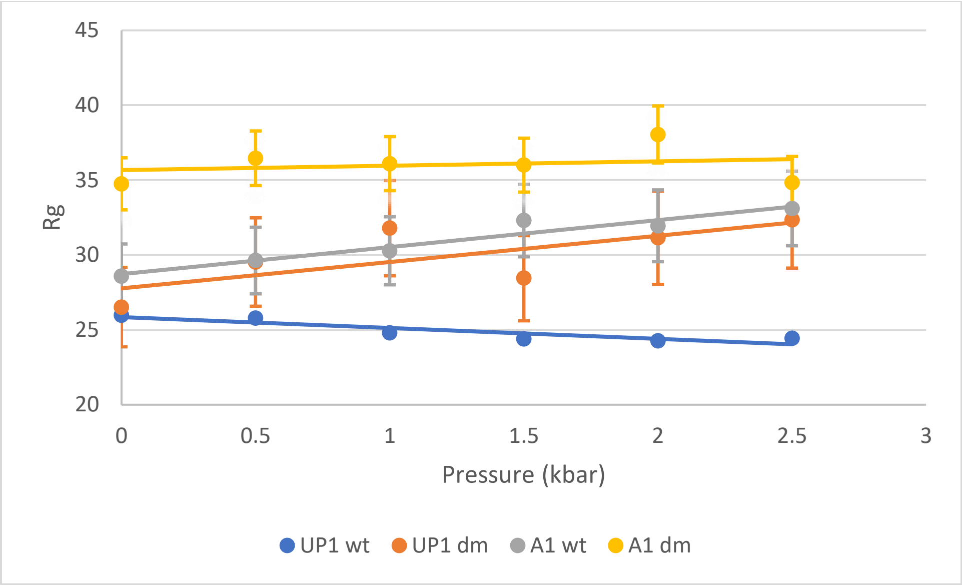


**Figure S8:** Numerical Rg values for UP1 and hnRNP A1 variants plotted versus pressure. Rg values were calculated from Guinier plots using BioXTAS RAW 2.1.4.

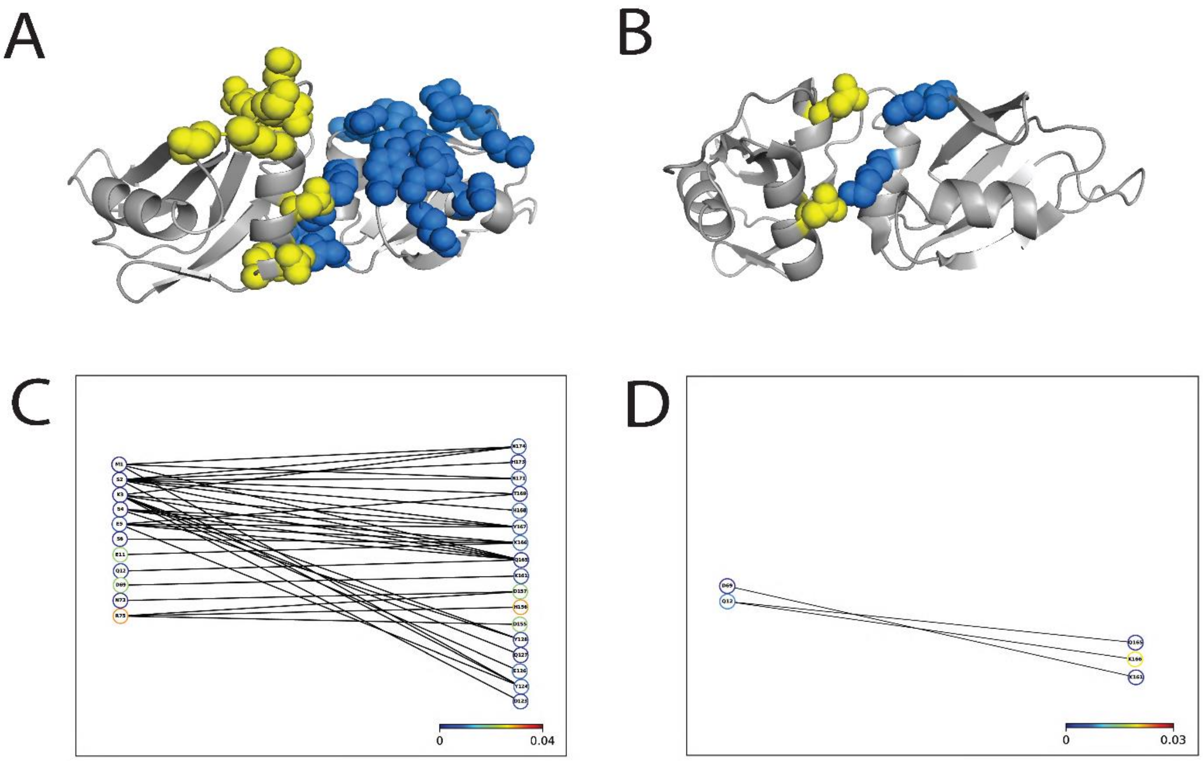

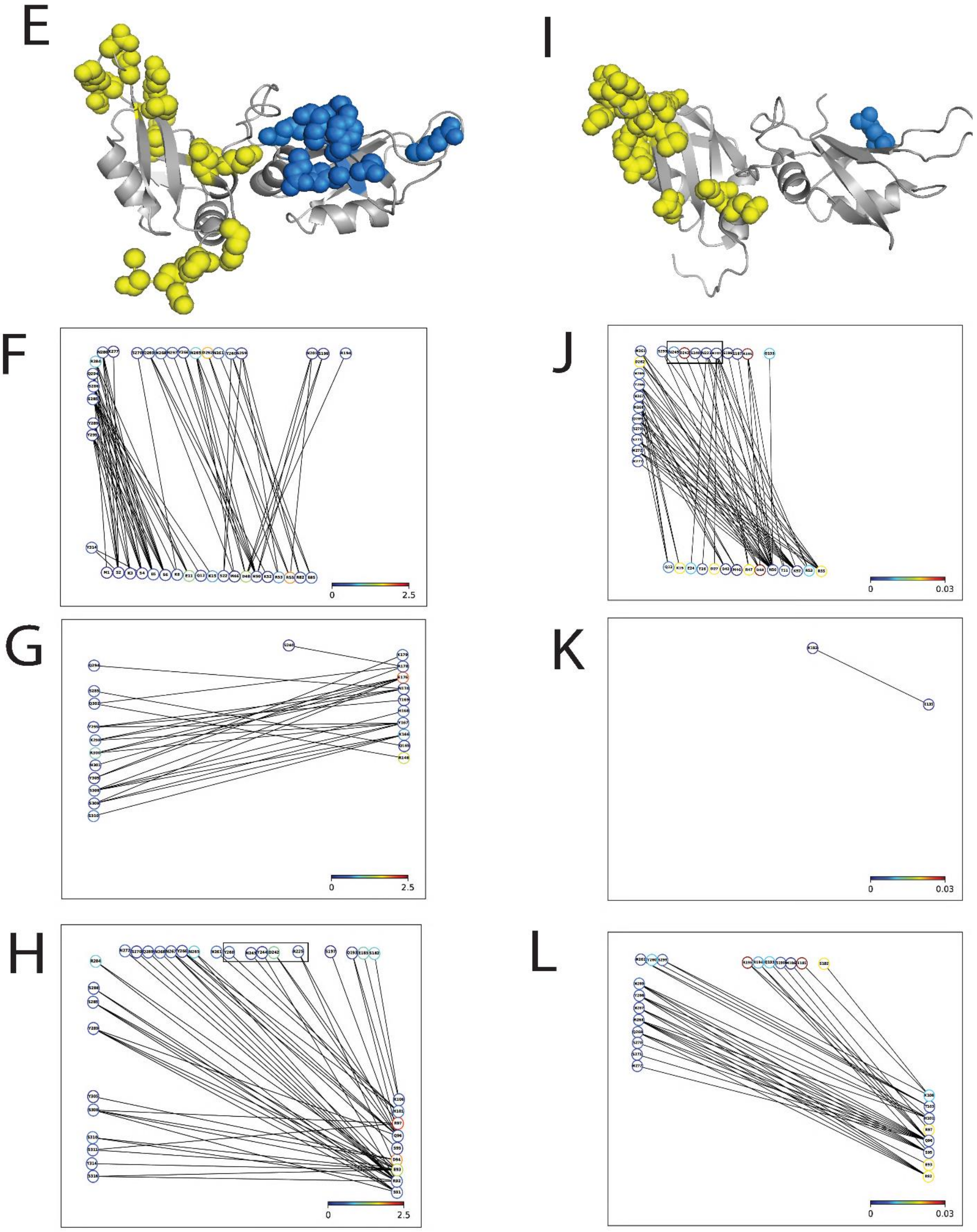
E
I
F
0
2.5
J
0
0.03
G
0
2.5
K
0
0.03
H
0
2.5
L
0
0.03

**Figure S9: Hbond network as determined by molecular dynamic simulations for UP1 and hnRNP A1 wt and dm variants A.)** WT inter-RRM mapped out onto UP1 B.) raw data for WT inter-RRM C.) $UP1^{dm}$ residues involved in Hbonds mapped onto UP1 D.) Raw data for $UP1^{dm}$ inter-RRM interactions E.) residues involved in UP1-$LCD_{A1}$ Hbond network for WT RRM1 and RRM2 F.) Raw data for WT RRM1-$LCD_{A1}$ interactions G.) Raw data for WT RRM2-$LCD_{A1}$ interactions H.) raw data for WT inter-RRM linker Hbond interactions with $LCD_{A1}$, RGG box residues are highlighted by the black box I.) $A1^{dm}$ residues involved in UP1-$LCD_{A1}$ interactions mapped out on RRM1 J.) raw data for $A1^{dm}$ RRM1-$LCD_{A1}$interactions, residues of the RGG box are highlighted by black box K.) Raw data for $A1^{dm}$ RRM2-$LCD_{A1}$ interactions, L.) raw data for $A1^{dm}$ inter-RRM interactions with $LCD_{A1.}$

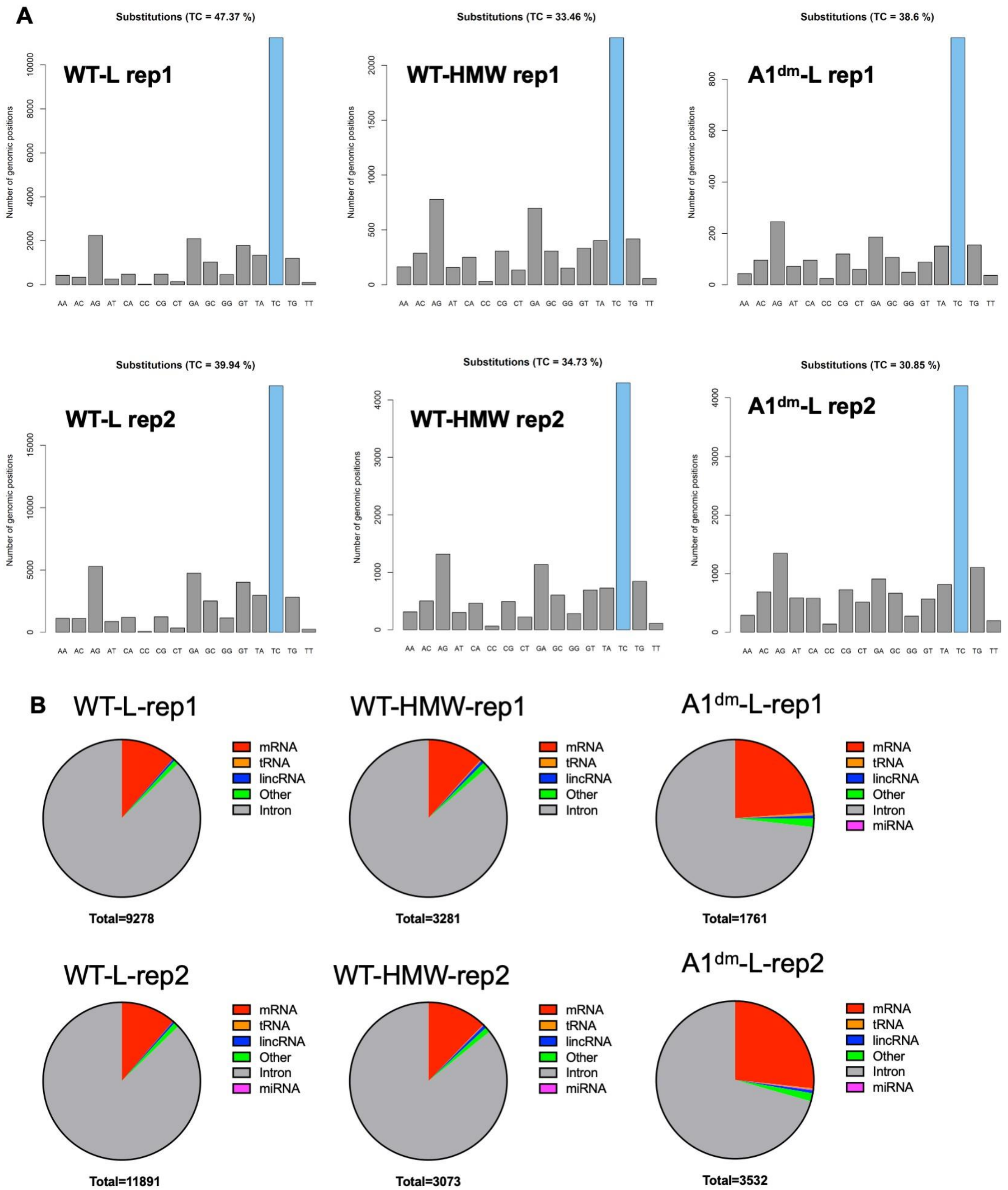

A
Substitutions (TC = 47.37 %)
WT-L rep1
Substitutions (TC = 33.46 %)
WT-HMW rep1
Substitutions (TC = 38.6 %)
A1dm-L rep1
Substitutions (TC = 39.94 %)
WT-L rep2
Substitutions (TC = 34.73 %)
WT-HMW rep2
Substitutions (TC = 30.85 %)
A1dm-L rep2
Number of genomic positions
AA AC AG AT CA CC CG CT GA GC GG GT TA TC TG TT
B
WT-L-rep1
Total=9278
WT-HMW-rep1
Total=3281
A1dm-L-rep1
Total=1761
WT-L-rep2
Total=11891
WT-HMW-rep2
Total=3073
A1dm-L-rep2
Total=3532
mRNA
tRNA
lincRNA
Other
Intron
miRNA

**Fig. S10: PAR-CLIP analysis of WT hnRNP A1 and the A1$^{dm}$ mutant. (**A) Frequency of the indicated nucleotide substitutions in PAR-CLIP-derived reads mapping to the human transcriptome are shown. (B) PAR-CLIP reads were formed into "clusters/binding sites" and annotated. Number of clusters that are present in the introns, mRNAs, tRNAs, lincRNAs, miRNAs and other RNA groups are shown.
